# *NPEPPS* segmental duplication drives position effect expression of *TBC1D3* in the human brain

**DOI:** 10.64898/2026.01.14.699559

**Authors:** Xavi Guitart, DongAhn Yoo, Jessie W Brunner, Hyeonsoo Jeong, Luyao Ren, Nidhi Koundinya, David Porubsky, Kendra Hoekzema, Katherine M Munson, Kaitlyn A Sun, Marcelo Ayllon, Kaylynn Hoglin, Reed McMullen, Bryan Pavlovic, Mitchell R Vollger, Alex A Pollen, Evan E Eichler

## Abstract

In humans, the *TBC1D3* gene family has been implicated in cortical expansion by promoting neuronal proliferation, yet the gene family exhibits extreme copy number variation (3-39 copies) and structural heterozygosity (97%). We investigate how a gene linked to development can remain so variable. We show that 45–96% of *TBC1D3* expression derives from a single paralog. The paralog’s >3-fold higher expression is driven by a ∼8.9 MYA segmental duplication that relocated a partial duplication of puromycin-sensitive aminopeptidase (*NPEPPS*), including its promoter, adjacent to *TBC1D3*. Using neurospheres and comparative transcriptomics of iPSC-derived cultures, we demonstrate that *NPEPPSP1–TBC1D3* expression increases during neuronal differentiation and *TBC1D3* is regulated by alternative splicing of a polyadenylation signal. Although the *NPEPPS* duplication is present in other apes, two separate structural events have compromised the gene fusion link in *Pan* and *Gorilla*, rendering this position effect human-specific.

## INTRODUCTION

*TBC1D3* is a primate-specific gene family that promotes cellular proliferation in both neurodevelopment and cancer. Originally identified in prostate and breast cancer tumors, functional experiments determined that *TBC1D3* promotes cell transformation and proliferation by manipulating cell vesicle transport, specifically amplifying the effect and duration of insulin-like growth factor (IGF) and epidermal growth factor (EGF) pathway-associated receptors (Frittoli et al., 2008, Wainszelbaum et al., 2008 & 2012). While its cell proliferation is co-opted in cancers, endogenous *TBC1D3* plays a developmental role by potentially expanding neural progenitor populations in the primate cerebral cortex, particularly in outer radial glia (Ju et al., 2016). Subcellular localization studies suggest that while *TBC1D3* is generally expressed cytoplasmically, in neuronal contexts *TBC1D3* translocates to the nucleus, where it inhibits the histone methyltransferase G9a (Hou et al., 2021). This repression reduces H3K9me2-mediated gene silencing and delays differentiation of outer radial glial cells, leading to additional division rounds and potentially contributing to the exponential increase in cortical neurons associated with human brain development.

The corresponding *TBC1D3* gene family is embedded in a dynamic core duplicon and has expanded recently during primate evolution via segmental duplications (SDs) across chromosome 17 (Jiang et al., 2007). While many of the orphan *TBC1D3* paralogs are thought to be nonfunctional pseudogenes, in humans, transcribed *TBC1D3* copies that maintain an open reading frame (ORF) originate primarily from two major clusters, Cluster 1 and Cluster 2, mapping to chromosome 17q12 (Guitart et al., 2024). These SD clusters also promote non-allelic homologous recombination leading to a recurrent 17q12 deletion/duplication (known as renal cyst and diabetes or RCAD syndrome) (Mitchel & Moreno-De-Luca et al., 1993; Mefford et al., 2007, 2016). Both the duplication and deletion syndromes manifest heterogeneously in patients, though the deletion syndrome is more often associated with abnormalities of the renal and endocrine systems. In contrast, the duplication syndrome is associated with both neurodevelopmental and psychiatric abnormalities. While 75% of deletion events are *de novo*, 90% of duplications are inherited, suggesting that the duplication may cause a less severe phenotype and thus persist in the general population for a few generations (Mitchel & Moreno-De-Luca et al., 1993; Mefford et al., 2016).

The origin and expansion of *TBC1D3* is complex: *TBC1D3* was proposed to originate from *USP6NL* (i.e., *RNTRE*)—given the conserved exon structure across 13 of 14 exons—though at the amino acid level the proteins share only 34% identity suggesting extensive amino acid replacement over a short period of primate evolution (Frittoli et al., 2008). In our prior work, we characterized the evolution and copy number diversity of the *TBC1D3* gene family among primates (Guitart et al., 2024). We showed *TBC1D3* expanded independently across seven separate lineages of the simian infraorder but is generally absent from prosimians. Within humans, *TBC1D3* is estimated to have expanded a human-specific modified version of the predicted protein ∼2 million years ago (MYA). This gene expansion coincides with the emergence of the *Homo erectus* (Bar-Yosef et al., 2001).

All human-expressed *TBC1D3* copies carry a human-derived 58 amino acid modification of the carboxy-terminus suggestive of a human-specific neofunctionalization (Guitart et al., 2024). Additionally, the gene family continues to dynamically mutate within the human population. Genome-wide studies have shown that *TBC1D3* is among the most copy number polymorphic and heterozygous gene families in the human species, ranging from 3-39 copies per haplotype genome (Sudmant et al., 2015; Guitart et al., 2024). Using long-read transcriptome sequencing to assign isoforms to specific paralogs, we showed that despite this variability, >90% of *TBC1D3* transcription from human fetal brain and induced pluripotent stem cells (iPSCs) maps to Cluster 2, and, more specifically, that over 80% of those transcripts stem from the last *TBC1D3* copy mapping distally at the edge of Cluster 2. We hypothesized that a position effect was responsible for regulating and restricting expression to the edge of Cluster 2.

To test this hypothesis and understand the nature of this potential position effect, we performed a detailed comparative transcriptomic and epigenetic analysis to investigate the regulation of *TBC1D3* in humans and great apes. We find that the Cluster 2 terminal *TBC1D3* paralog dominates expression as the result of fusion with an upstream duplicated gene, *NPEPPSP1*, including its brain-enriched promoter. This *NPEPPS1-TBC1D3* fusion transcript accounts for the majority of *TBC1D3* expression across various tissues. We show that the fusion is mediated by segmental duplication of *NPEPPS* in the common ancestor of African great apes but, by two independent events, was lost in the chimpanzee, bonobo, and gorilla lineages by subsequent gene conversion and deletion events, respectively. Additionally, while both *NPEPPSP1* and *TBC1D3* are copy number polymorphic in humans, we find that the promoter of the fusion gene is fixed in all human genomes examined. Our findings dissect the evolution of the *TBC1D3* position effect and explain how it has become specific to the human lineage providing a model for how new genes rapidly emerge in a species.

## RESULTS

### Matched *TBC1D3* copy expression and regulation in CHM13

The high sequence identity (>99%) of *TBC1D3* paralogs has meant that the transcription and regulation of this gene family has essentially been excluded from large-scale studies such as ENCODE (ENCODE Project Consortium, 2012). To resolve this, we initially used matched long-read sequencing (LRS) data from the complete human reference genome (T2T-CHM13) to investigate the methylation profile of the nine *TBC1D3* copies present in this haploid source genome (Fig. 1A; Nurk et al., 2022; Methods). This analysis revealed a distinct hypermethylation signature across the gene body of the terminal paralog of Cluster 2, *TBC1D3-CDKL2*, whereas all other *TBC1D3* paralogs displayed uniformly low methylation signals, comparable to background levels observed for lowly expressed genes, including SD paralogs (Vollger et al., 2022). This hypermethylation signature of the *TBC1D3-CDKL2* gene body aligns with one of the canonical epigenetic hallmarks of expressed genes.

**Figure 1.**
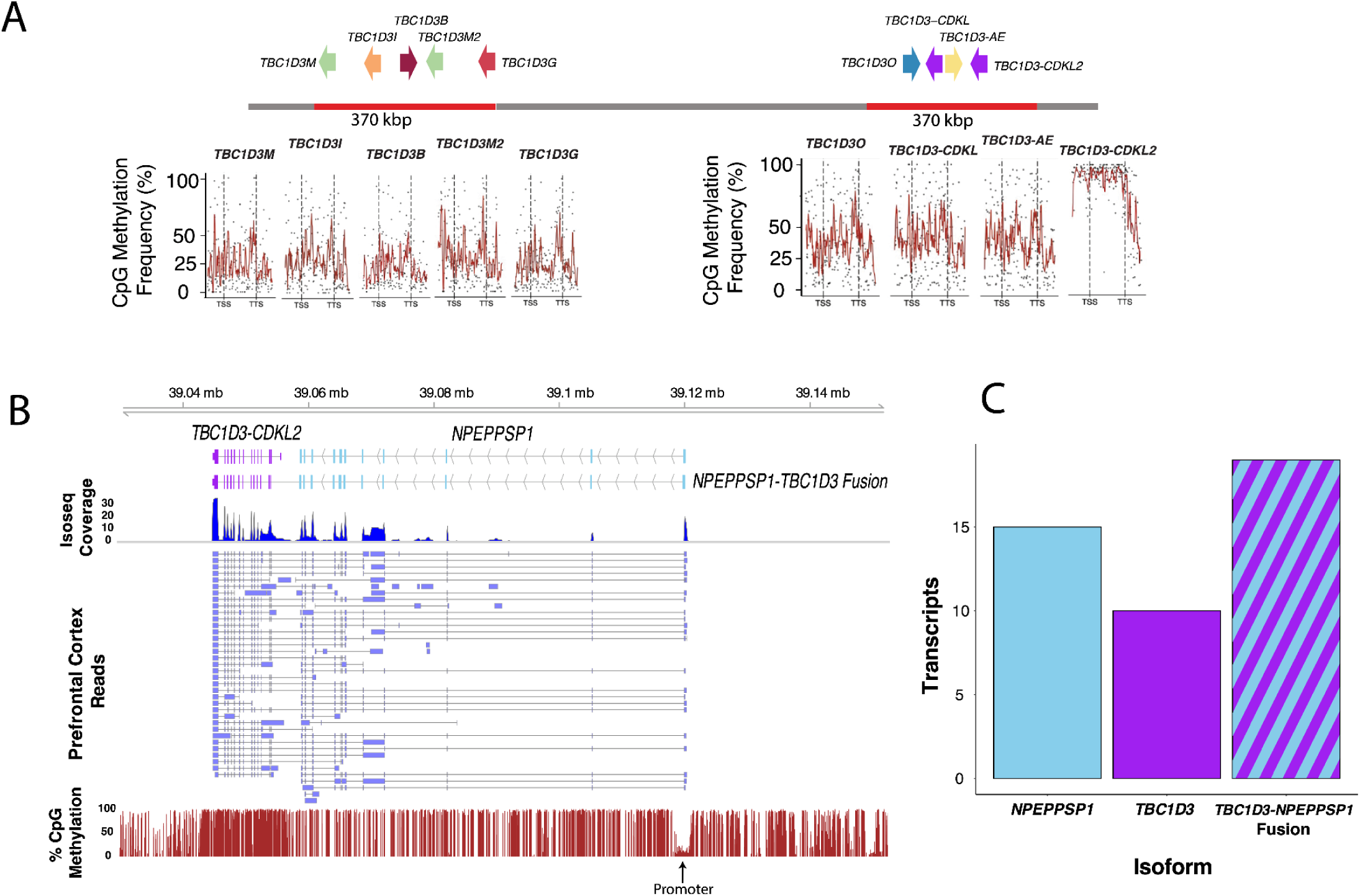
Differential regulation and expression of the human *TBC1D3* gene family. **A.** CpG methylation profile compared among *TBC1D3* paralogs within CHM13 based on mapping of CHM13 ONT data back to the T2T-CHM13 genome (Nurk et al., 2022). Percent methylation of CpG sites from transcription start site (TSS) to transcription termination site (TTS) based on rolling average of 15 consecutive CpG sites (red lines). This methylation signature is consistent with >75% of transcription originating from this distal copy (Guitart et al., 2025). **B.** Expression and promoter definition of the terminal *TBC1D3-CDKL2* copy. A schematic of the *NPEPPSP1-TBC1D3* fusion gene structure, including overall Iso-Seq transcript coverage per exon (top panel) and various isoforms (middle panel) detected based on Iso-Seq transcript data from human prefrontal cortex (Leung et al., 2021). Percent CpG methylation data from CHM13 overlaid with this annotation predicts the likely location of the promoter identified by the dip in methylation (bottom panel). **C.** The absolute number of Iso-Seq transcripts identified as *NPEPPSP1-TBC1D3* fusion vs. solo *TBC1D3* or *NPEPPS1* compared for Iso-Seq data from CHM13.

To identify the putative promoter driving expression of this distal *TBC1D3-CDKL2* copy, we analyzed the flanking 100 kbp region upstream of its transcription start site (TSS) for evidence of transcription initiation. An analysis of short-read ATAC-seq data and annotated ENCODE cis-regulatory elements in the GRCh38 reference suggested a putative open chromatin accessibility region located 66 kbp from the translation initiation site of *TBC1D3* (Supplementary Fig. 1; ENCODE Project Consortium, 2012). To define the full-length gene structure based on empirical data rather than annotation alone, we mapped full-length Iso-Seq transcriptomic data from human prefrontal cortex (Fig. 1B). Remarkably, 43% (19/44) of transcripts did not align to the canonical RefSeq model but instead represented a fusion isoform with an annotated pseudogene, *NPEPPSP1*, located directly upstream (Fig. 1C). Tracing this dominant isoform to its TSS revealed a distinct dip in methylation immediately upstream of the first exon—a clear hypomethylated promoter signature, consistent with active transcription initiation (Gershman et al., 2022; Vollger et al., 2022).

### Fusion expression and regulation in *in vitro* developing brain model

While CHM13 proved useful to discover the potential site of transcription initiation, this developmental abnormality—a hydatidiform mole—is a poor proxy for the human developing brain, the principal tissue of *TBC1D3* function (Ju et al., 2016; Hou et al., 2021). To investigate *TBC1D3* regulation in a context more relevant to brain development, we first interrogated human neurospheres, an *in vitro* model for neural differentiation harboring both neural progenitor cells (NPCs) and neural stem cells (Fig. 2A; Real et al., 2025). We profiled both chromatin accessibility and transcriptome activity of the *TBC1D3* gene family in neurospheres derived from the HPRC sample HG02630.

**Figure 2.**
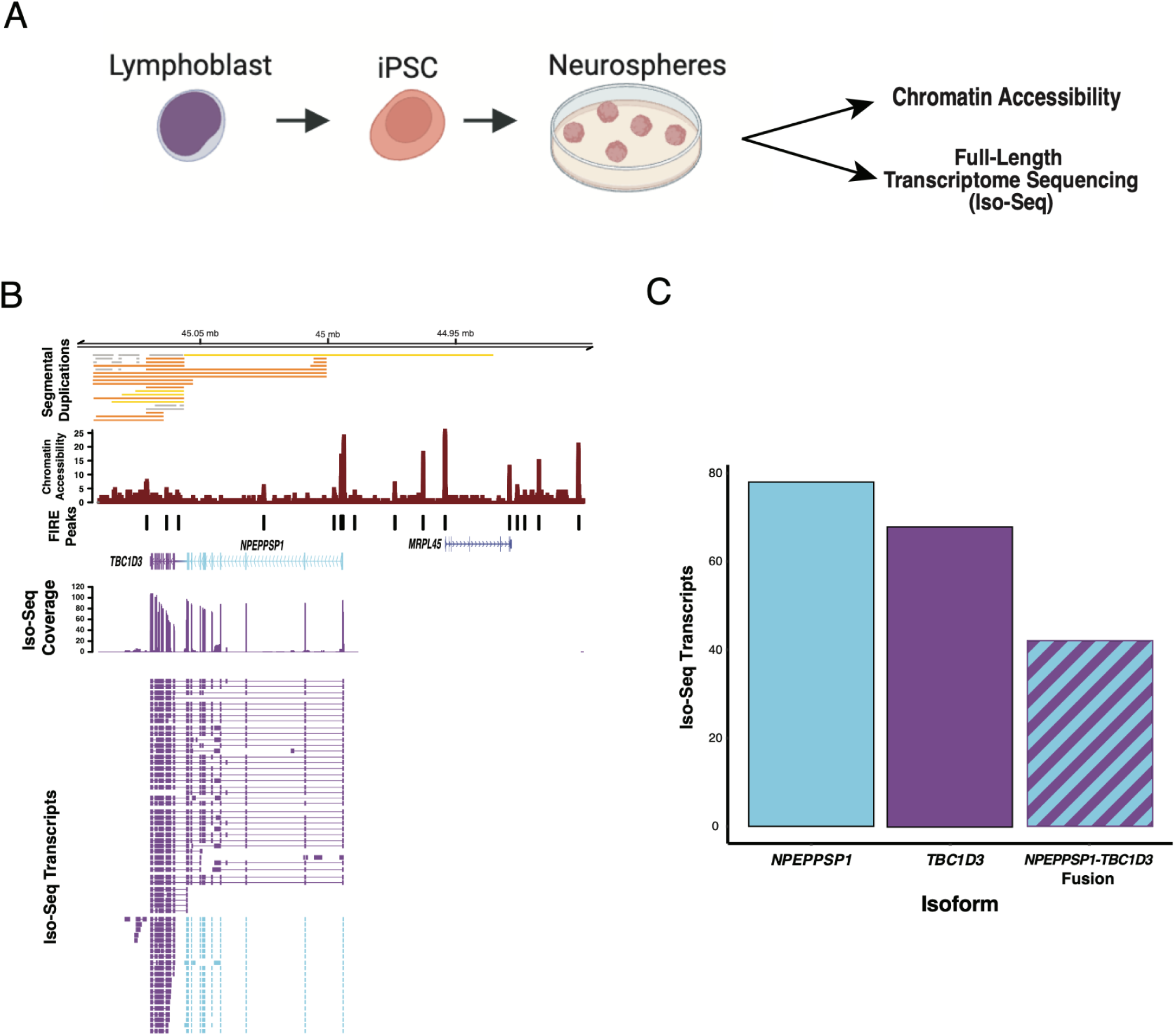
Transcription and chromatin accessibility in human neurospheres. **A.** Schematic depicting neurosphere generation. Lymphoblasts from the 1000 Genomes Project (1KGP) sample, HG02630, were induced to create iPSCs and subsequent neurospheres. Matched Iso-Seq and Fiber-seq data were generated from neurospheres to assess the transcriptome and chromatin accessibility. **B.** Expression and chromatin accessibility of *NPEPPSP1-TBC1D3*. Full-length transcripts mapped to the HG02630 diploid *de novo* sequence assembly are shown with alignment coverage, Fiber-seq accessibility, and segmental duplication sequence annotated with SEDEF (Numanagic et al., 2018). **C.** Absolute expression of *NPEPPSP1*, *TBC1D3*, and the *NPEPPSP1-TBC1D3* fusion. Transcript reads are classified into *NPEPPSP1*, *TBC1D3*, and *NPEPPSP1-TBC1D3* fusion categories.

Consistent with CHM13 methylation data, this analysis revealed that none of the *TBC1D3* paralogs showed evidence of a proximal promoter—defined as a peak of accessibility directly upstream of the TSS of the canonical gene model (Supplementary Table 1; Stergachis et al., 2020). Instead, the matched transcriptomic-DNA genome assembly data derived from the same neurosphere source material confirmed the initial observation from the analysis of the prefrontal cortex library; 56% (116/206) of primary *TBC1D3* transcript mappings were transcribed as a fusion product with *NPEPPSP1*, over three times the expression than any other individual *TBC1D3* paralog. Using Fiber-seq Inferred Regulatory Elements (FIRE) (Fig. 2B; Vollger et al., 2025), we defined the promoter element at the *NPEPPSP1* TSS, a 302 bp unit that regulates expression of the *TBC1D3* paralog located directly downstream of *NPEPPSP1*. Combined, these data suggest that cooption of the *NPEPPSP1* promoter plays a significant role in generating neuronal transcripts of *TBC1D3*, especially from the most distal copy of the *TBC1D3* Cluster 2 (Fig. 2B,C).

### Juxtaposition of novel regulatory DNA by segmental duplication of NPEPPS

To understand the evolutionary origin of the regulatory sequence driving *NPEPPSP1-TBC1D3* expression, we examined its genomic context in the HG02630 diploid assembly. Using SD annotations by SEDEF (Numanagic et al., 2018), the *NPEPPS1* duplication and its associated promoter arose as a 119,358 bp SD originating from *NPEPPS* located 9.16 Mbp distally on chromosome 17q21 (Fig. 3). A comparison of these two paralogous regions along with corresponding gene annotation of the ancestral locus shows that the promoter sequence of *NPEPPS* (N-peptide puromycin sensitive) matches (Fig. 3A) the promoter sequence associated with *NPEPPS1-TBC1D3* fusion transcripts defined by LRS (Fig. 2). The ancestral *NPEPP*S encodes a zinc metallopeptidase broadly expressed across tissues, and mutations of it have been associated with neurodegenerative tauopathies and Parkinsonism (Henderson Front Gent 2021; Karsten et al., 2006). Thus, the two paralogous promoter pairs regulate expression of *NPEPPS* and *NPEPPSP1*, and *MRPL45* and *MRPL5P1*, respectively (Fig. 3A). The duplicated region shows an average of 98.2% sequence identity by matches only, suggesting a recent origin during ape evolution.

**Figure 3.**
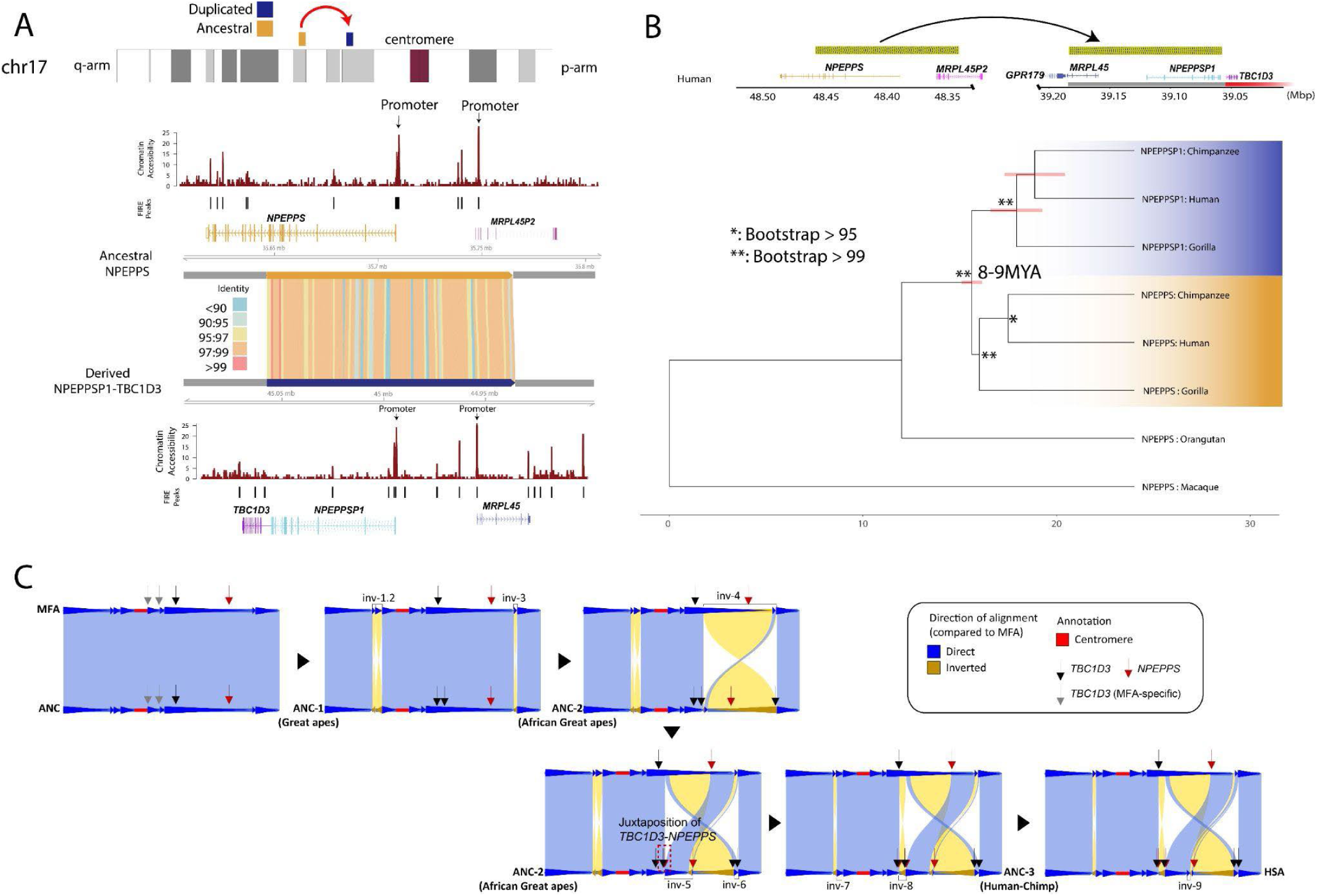
Evolutionary origin of *NPEPPSP1* regulatory sequence. **A.** Comparison of the sequence and regulatory landscape of *NPEPPS* and *NPEPPSP1*. SVbyEye reveals a ∼119 kbp SD with 98.2% sequence identity between the ancestral *NPEPPS* and duplicated *NPEPPSP1* loci. Fiber-seq data analyzed using FIRE compared chromatin accessibility for *NPEPPS* (above) to *NPEPPS1-TBC1D3* (below) with the most likely site of the promoter (indicated). **B.** *NPEPPS1* segmental duplication evolution. A phylogeny of 15 kbp of the *NPEPPS* duplication most proximal to *TBC1D3* and macaque (25 MYA divergence) predicts that the duplication occurred ∼8-9 MYA. **C.** Based on telomere-to-telomere (T2T) comparative chromosomal analyses and assuming the macaque genome represents the ancestral archetype, we develop a parsimonious model to estimate the timing and progression of the eight ancestral and four lineage-specific inversions during ape evolution (Supplementary Fig. 4). The top track represents macaque while bottom track represents the predicted intermediate states during ape evolution.

In order to more accurately estimate the timing of the duplication, we extracted a 15 kbp region from the *NPEPPSP1*-anchored end of the SD, directly adjacent to *TBC1D3*. We used this sequence to identify homologous sequences in human, chimpanzee, gorilla, orangutan, and macaque genome assemblies (Yoo et al., 2025). Our analysis shows that the ∼119 kbp *NPEPPS* SD is duplicated in all African apes assessed (human, chimpanzee, bonobo, and gorilla) (Fig. 3; Supplementary Figs. 2, 3) but is present only as a single unique locus in orangutan and macaque, corresponding to *NPEPPS*. This confirms *NPEPPS* as the ancestral locus and that the duplication (*NPEPPSP1*) arose in the common ancestor of all African great apes. We also constructed a maximum likelihood phylogenetic tree by generating a multiple sequence alignment of all copies among humans and apes using macaque as an outgroup (Fig. 3B). We estimate that the duplication occurred in the common ancestor of African great apes approximately 8.49 MYA (95% confidence interval of 8.00–9.42 MYA) (Fig. 3B).

A comparative analysis of human chromosome 17 among primate genomes reveals that both *TBC1D3* genes as well as *NPEPPS1-TBC1D3* are associated with series of large-scale nested inversions changing the gene order of over 90 Mbp of this chromosome during primate evolution. As noted previously (Guitart et al., 2024), *TBC1D3* often maps at or near the breakpoints of these large-scale evolutionary rearrangements (Supplementary Figs. 4, 5). Based on the completed sequence of macaque and six other nonhuman primates (Yoo et al., 2025), we developed a parsimonious model to account for the series of inversion events (Fig. 3C) that likely restructured this region over the course of evolution and predict 12 large-scale evolutionary inversions (at least 8 ancestral and 4 species-specific events). This includes three African great ape ancestral inversions (inv-4-6), a human–chimpanzee ancestral inversion (inv-8), and a human-specific inversion (inv-9); these five inversions are directly flanked by *TBC1D3* or *NPEPPS* genes. All inversions range in size from 939 kbp to 44 Mbp (Supplementary Table 2). In the case of the *Pan* lineage, the pericentric inversion relocates both the *TBC1D3* clusters and *NPEPPS1-TBC1D3* from the q arm to the p arm. Focusing specifically on the terminal *TBC1D3-NPEPPSP1*, our analysis suggests that the duplication of *NPEPPSP1* to *TBC1D3* occurred in a common African ape ancestor (inv-5) in conjunction with large-scale inversions of that region in the ancestral lineage (Fig. 3C; Supplementary Fig. 5). Notably, examination of additional human haplotypes from the Human Pangenome Reference Consortium (HPRC) (Lucas J. K. et al. 2026) revealed that the ancestral nonhuman ape orientation is still present, including inv-7, inv-8, and inv-9, although at a low allele frequency (2.8, 0.6, and 9.2%, respectively; Supplementary Table 6; Supplementary Fig. 6). This may suggest longstanding recurrent inversions or evolutionary toggling (Porubsky et al., 2020) and that the restructuring of the locus is still ongoing among humans.

### Comparative transcriptomics in differentiating neurons

To investigate how *NPEPPSP1-TBC1D3* fusion expression changes across neuronal development, and to compare this regulation between humans and other apes, we designed a three-stage cell culture experiment using fibroblast-derived iPSCs from human, chimpanzee, and orangutan (the latter serving as an outgroup; Fig. 4A). Cells were induced sequentially from iPSCs into NPCs, and finally mature neurons with the identity confirmed by marker analyses (Jeong et al., in preparation). At each stage, we collected both genomic DNA (gDNA) and full-length transcriptomes using HiFi sequencing and Kinnex Iso-Seq, respectively. We also generated a donor-specific genome assembly (DSA) for each sample using iPSC gDNA and included methylation calling during base calling to track regulatory changes throughout development. This resource allowed us to match transcript, methylation, and genomic data to specific paralogs without reference bias or cross-mapping among high-identity paralogs.

**Figure 4.**
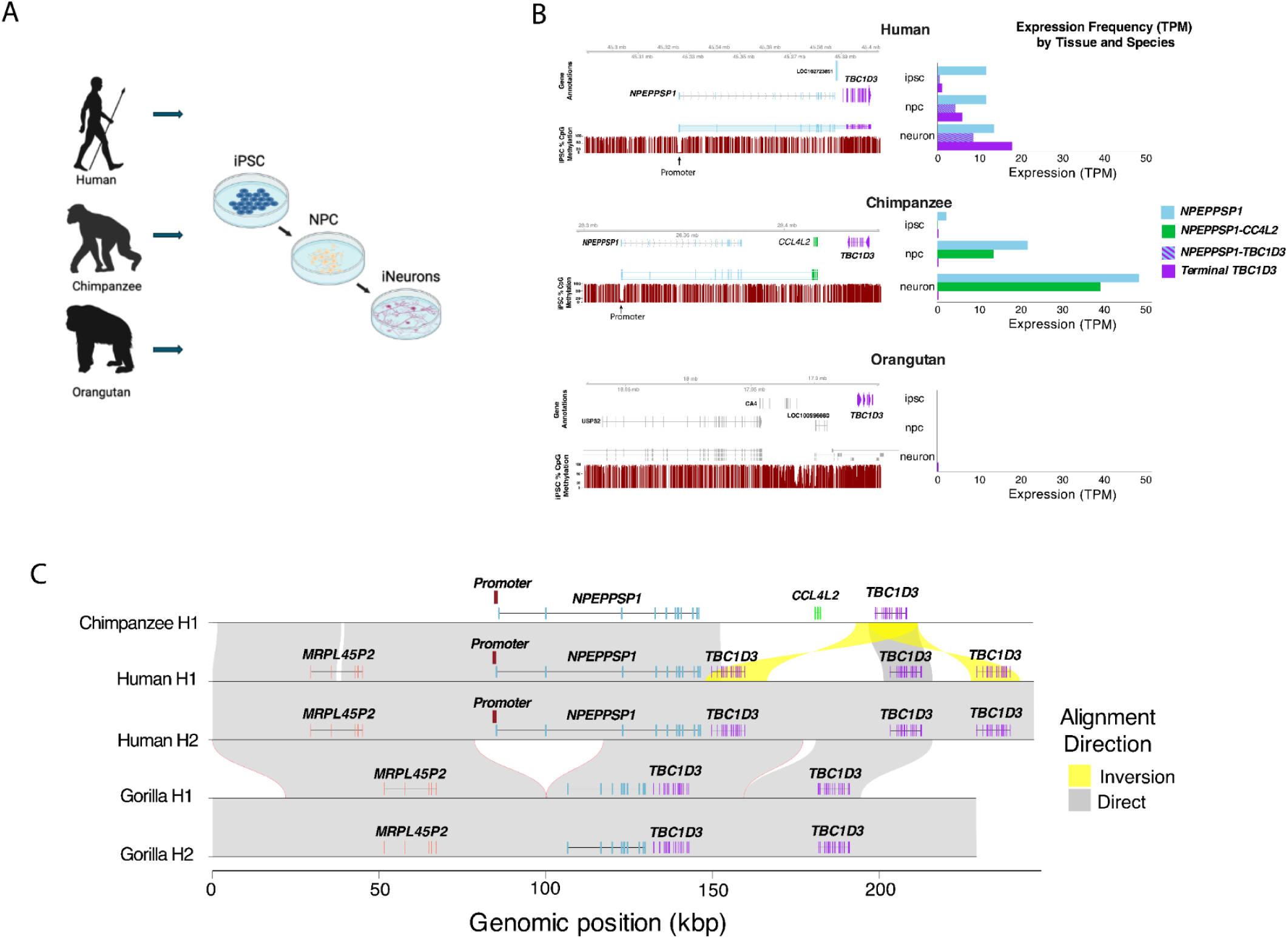
Comparative expression of *NPEPPSP1-TBC1D3* in a neuronal developmental cell culture model of great apes. **A.** Experimental design. Fibroblasts sourced from a human, chimpanzee, and gorilla samples were induced into iPSCs, NPCs, and neurons. Donor-specific genome assemblies from the iPSCs as well as gDNA and mRNA were generated using long-read sequencing protocols for each of the three stages. **B.** Comparative methylation and transcription of *NPEPPSP1*, terminal *TBC1D3*, and fusion genes in human, chimpanzee, and orangutan. A schematic of the gene organization and methylation profile (red; % of read methylated over CpG) is shown for iPSCs from each of the three species (left panels). Expression levels (transcripts per million, TPM) are compared for *NPEPPS1*, fusion genes, and the terminal *TBC1D3* copy for each of three stages (right panels). Histogram illustrates TPM based on Kinnex sequencing (Methods). **C.** Alignments of the *NPEPPSP1* duplication locus between one chimpanzee, two human, and two gorilla haplotypes show a 38 kbp deletion fixed in the gorilla lineage, removing both the *NPEPPSP1* promoter and first two exons.

In humans, we observe stable *NPEPPSP1* expression across all three proxy developmental stages, accompanied by a stepwise increase in *TBC1D3* expression from iPSCs to mature neurons (Fig. 4B). This increase of *TBC1D3* expression was matched by a correlated increase in the *NPEPPSP1-TBC1D3* fusion transcript levels, increasing from a single identifiable fusion transcript in iPSCs to 72% (18/25) and 47% (98/208) of *TBC1D3* expression in NPCs and neurons, respectively. As expected, the fusion transcript was driven by the predicted juxtaposed promoter, characterized by the methylation dip at the *NPEPPSP1* TSS, consistent with the FIRE-defined regulatory elements observed in HG02630 (Fig. 3A; Vollger et al., 2025).

In contrast, chimpanzee *TBC1D3* expression is minimal throughout our model of neuronal differentiation. While both lineages show activation as well as a stepwise increase of the same orthologous *NPEPPSP1* promoter as cells differentiate to neurons, in chimpanzees that stepwise increase in *NPEPPSP1* expression across development is not associated with *TBC1D3* but rather a different gene, *CCL4L2* (CC chemokine ligand 4), a protein associated with inflammation and immune response (Li et al., 2023; Xu et al., 2024). Two key observations help explain the absence of *NPEPPS1-TBC1D3* expression in chimpanzee. First, the downstream *TBC1D3* paralog in the chimpanzee haplotype is inverted with respect to its ORF relative to the *NPEPPSP1* promoter, precluding *NPEPPSP1-TBC1D3* gene expression. Second, we identified a 44 kbp insertion of *CCL4L2* situated between the *NPEPPS1* promoter and *TBC1D3*, resulting in abundant *NPEPPSP1–CCL4L2* fusion transcription in chimpanzees (Fig. 4B). Thus, in chimpanzee, *CCL4L2* appears to co-opt the *NPEPPSP1* promoter analogous to human *TBC1D3* but with a very different outcome. Analysis of the bonobo (*Pan paniscus*) genome indicates that both the inversion and *CCL4L2* insertion are present in this species, suggesting that this organization is a longstanding property of the *Pan* genus likely originating in the common ancestor of the two extant lineages (Supplementary Figs. 2, 3).

Orangutan, which does not harbor the *NPEPPS1* SD or its promoter, serves as a negative control and shows scant *TBC1D3* expression of the most terminal copy (Fig. 4B). We note, however, that there is abundant *TBC1D3* expression in the orangutan (Supplementary Fig. 7) from other copies. However, unlike the human ortholog, orangutan *TBC1D3* is uniformly expressed by numerous internal Cluster 2 paralogs that have independently expanded in the *Pongo* genus (Guitart et al., 2024; Supplementary Fig. 8). While gorilla was not included in our neuron development cell model, we also attempted to investigate *NPEPPSP1-TBC1D3* regulation by exploring the orthologous locus in the gorilla genome assembly (Yoo et al., 2025). While gorilla inherited the same *NPEPPSP1* duplication, the promoter and first two exons of the pseudogene have been ablated by a subsequent 38 kbp deletion in the gorilla lineage (Fig. 4, Supplementary Fig. 3). Concomitantly, we find no evidence of *NPEPPSP1* promoter-initiated *NPEPPS1-TBC1D3* fusion transcripts in this species.

### The fusion transcript encodes two separate ORFs

Based on the human *NPEPPSP1-TBC1D3* fusion transcript consensus sequence, we predict two large, mutually exclusive ORFs (Supplementary Fig. 9A). The first ORF (ORF1) comprises a 449-codon reading frame corresponding to the entirety of *NPEPPSP1* and four codons located at the beginning of *TBC1D3* exon 2, traversing the fusion junction. The second ORF begins 13 bases upstream of ORF1, also at the second exon of *TBC1D3*, and represents the canonical 546-aa TBC1D3 ORF annotated by RefSeq (O’Leary et al., 2015). These ORFs are offset by two bases, placing them in different frames, precluding the formation of a fused protein product.

Unlike *TBC1D3*, there has been no characterization of *NPEPPSP1*, a supposed pseudogene, at the protein level. We searched publicly available proteomic resources using an *in silico* tryptic digest of *NPEPPSP1-TBC1D3* ORF1 and found four uniquely identifying peptides distinguishing *NPEPPSP1* from any other protein, including its ancestral paralog, *NPEPPS*. We searched for these four peptides in publicly available mass spectra datasets and identified five proteomic experiments with matches to the peptides. Although not conclusive, these data suggest *NPEPPSP1* may be translated and present in the human proteome (Supplementary Table 3; Methods).

Next, we assessed mutational tolerance of both ORFs by analyzing 295 human genomes recently sequenced as part of the HPRC (Liao et al., 2023) and the Human Genome Structural Variation Consortium (HGSVC) (Logsdon et al., 2025). Across these haplotypes, we found three instances of a premature termination of *NPEPPSP1*, driven by one splice junction in the third exon, and two nonsense mutations on the seventh exon (Supplementary Fig. 9B). By contrast, we found a single occurrence of a splice junction mutation (3’ ss of exon 10) introducing a premature stop codon in the *TBC1D3* ORF (Supplementary Fig. 9C).

### Posttranscriptional processing of fusion transcripts

Because *NPEPPSP1* duplicates only the first 11 exons of *NPEPPS*, we were interested in understanding how this partial duplication impacts the transcription and splicing of the gene. Using our neuronal developmental dataset, we characterized the expression of the two most common isoforms. We observe that solo *NPEPPSP1* transcripts utilize a novel polyadenylation site located 111 bp downstream of the canonical splice junction used by *NPEPPS* (Fig. 5A). This alternative site is strongly favored in iPSCs, being used in >98% (51/52) of full-length transcripts sequenced. The equivalent paralogous poly-adenylation site, however, was not observed in the ancestral *NPEPPS* gene and is excised within the 11th intron by a splice junction upstream of the site. Of note, we find that as cells differentiate toward neurons, *NPEPPSP1* splice site usage shifts toward the equivalent splice junction used by *NPEPPS*, losing the premature polyadenylation site and extending transcription to include the full-length *TBC1D3* ORF (Fig. 5B). Our results suggest that the fusion is regulated posttranscriptionally by alternative splicing of the last and first exons of *NPEPPSP1* and *TBC1D3*, respectively (Fig. 5).

**Figure 5.**
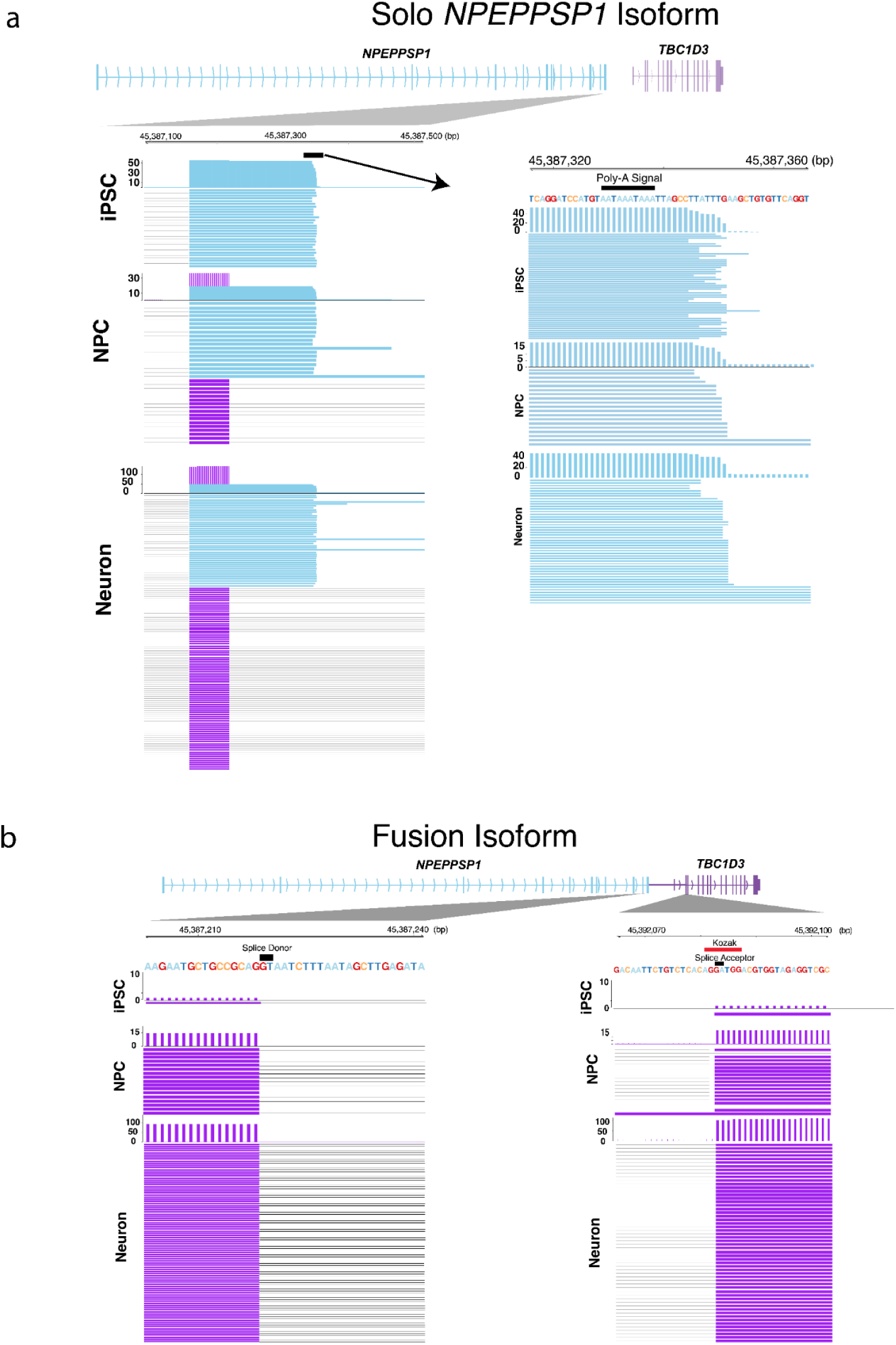
Alternative splicing of *NPEPPSP1-TBC1D3* fusion. **A.** Solo *NPEPPSP1* isoform. Solo *NPEPPSP1* transcripts display a polyadenylation signal (chr17:46387329-46387338) terminating the transcript, a signal that is removed by splicing in fusion genes. **B.** Fusion isoform. *NPEPPSP1-TBC1D3* transcripts display splicing between *NPEPPSP1* exon 10 and *TBC1D3* exon 2, where ORF2 begins. The splice acceptor site of *TBC1D3* overlaps with a Kozak sequence, recognized as AXXATGG.

We next sought to understand how the fusion impacts transcript processing. Given that nonsense-mediated decay (NMD) is often triggered by splice junctions located downstream of a translation stop codon, we hypothesized that *TBC1D3*-containing transcripts might be targeted for degradation and, thus, preferentially rescued upon NMD inhibition. To test this, we treated human and chimpanzee iPSCs and NPCs with cycloheximide (a known NMD inhibitor) or dimethyl sulfoxide (DMSO) as a control (Cheng et al., 2025). While we observe an overall increase in expression of *NPEPPSP1*, *TBC1D3*, and the fusion transcript following cycloheximide treatment, we did not observe preferential rescue of *TBC1D3*. Instead, *NPEPPSP1* transcripts show the most pronounced increase (Supplementary Fig. 10), suggesting that they may be more susceptible to a translation-dependent RNA degradation than *NPEPPS1-TBC1D3* fusion transcripts.

Importantly, we find that *TBC1D3* expression surpasses *NPEPPSP1* in transcript abundance as human cells differentiate into neurons (Supplementary Figs. 11–13; Supplementary Tables 4, 5). Initially, we considered the possibility that this was due to expression from other *TBC1D3* paralogs. However, both epigenetic profiling and full-length transcript analysis across paralogs rule out this explanation. All 196 transcripts map to the terminal Cluster 2 *TBC1D3* copy. Only a single hypomethylation signature is identified corresponding to the promoter of the fusion transcript. We fail to identify any methylation or chromatin accessibility signatures consistent with a promoter in another paralog (Supplementary Tables 1, 6).

While it is possible that this neuronal enrichment of *TBC1D3* may be due to the moderate 3’ bias of the Kinnex and Iso-Seq LRS platforms, this seems unlikely since it is not observed in either NPCs or iPSCs, which were processed identically with respect to capture. We speculate, therefore, that there may be a posttranscriptional mechanism that isolates the downstream *TBC1D3* reading frame from its upstream linked partner, *NPEPPSP1*. In support of this, we identified a Kozak sequence, AXXATGG, in the second exon of *TBC1D3*, located at the acceptor site of the splice junction fusing both genes, and including the start coding of ORF2, and suggest this site serves as the site initiating translation of the *TBC1D3* ORF (Fig. 5B; Kozak, 1984).

### Human population genetic analyses and structural variation

We extended our previous analysis of human variation (Guitart et al., 2024) by analyzing an additional 272 genomes recently released as part of the HPRC, HGSVC, and Platinum Genomes Project (Liao et al., 2023; Ebert et al., 2021; Porubsky et al., 2025). Of the 632 sample haplotypes, along with reference assemblies of T2T-CHM13 and GRCh38, 337 assemblies passed quality control (Methods). This updated analysis confirms that the two *TBC1D3* clusters are among the most structurally heterozygous gene families in the human species, with an overall estimated structural heterozygosity of ∼97%. Cluster 2, which is associated with the juxtaposed *NPEPPSP1* promoter, exhibits a slightly narrower distribution of *TBC1D3* copy number compared with Cluster 1 (range 2–14 vs. 1–22 copies; Fig. 6A) and a modestly lower mean copy number (Cluster 2 mean = 6.1; Cluster 1 mean = 6.9; *p* = 0.0004, Welch’s *t*-test, Supplementary Fig. 14). Concomitantly, Cluster 1 shows a 1.44 Mbp or a 6.5-fold length difference between the longest and shortest human haplotype (Fig. 6A) Within both clusters, however, *TBC1D3* copy number is significantly higher among individuals of African ancestry than among non-African samples (Cluster 1: African = 8.3 copies, non-African = 5.9, *p* = 8.4E-14; Cluster 2: African = 7.8, non-African = 5.2, *p*=8.2E-16; Welch’s *t*-test; Supplementary Fig. 14). Despite this incredible diversity, the *NPEPPSP1* promoter itself remains constrained, both in terms of structure and copy number (copy 4 in diploid genomes; Supplementary Fig. 15).

**Figure 6.**
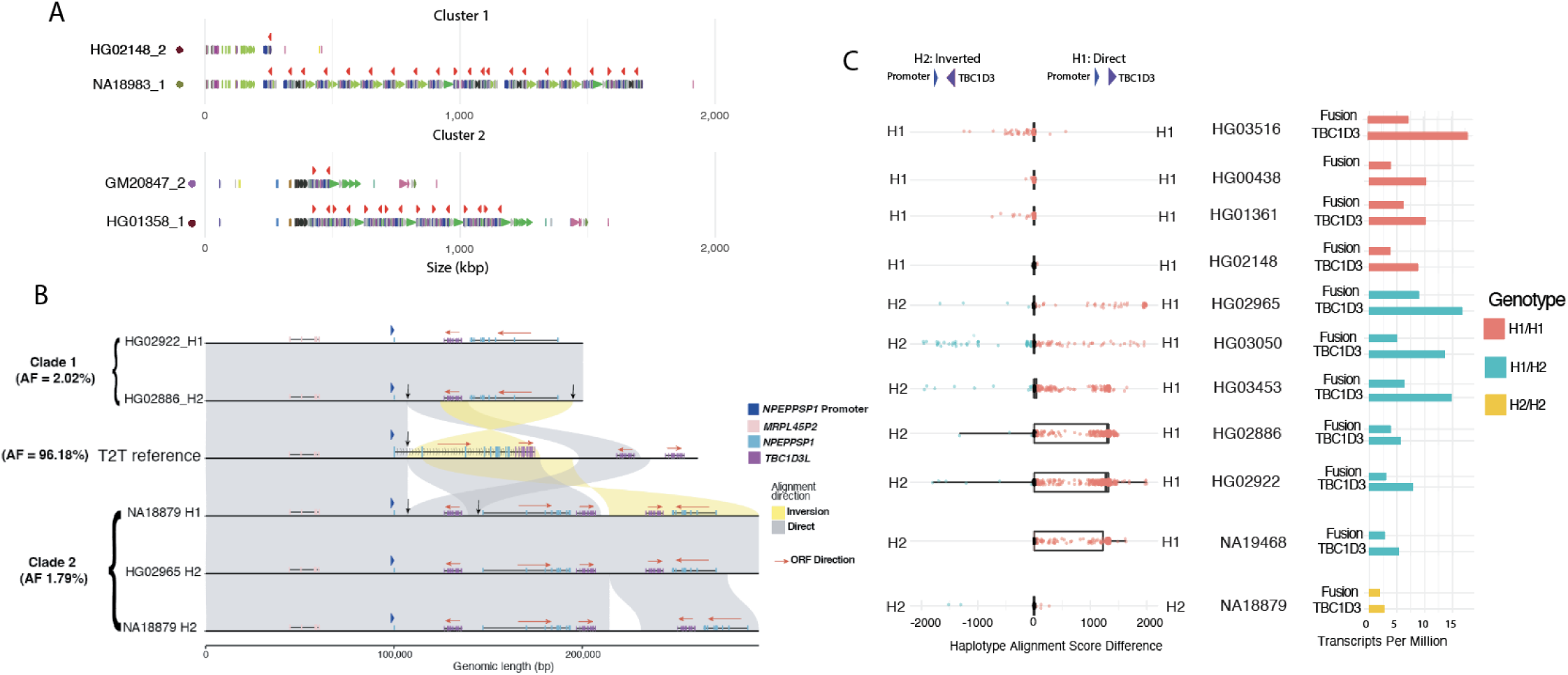
Human variation and inversion haplotypes. **A.** Cluster minimal and maximal haplotypes. Cluster 1 and 2 haplotypes vary between 1-22 and 2-15 copies, respectively. The largest and smallest haplotypes with respect to *TBC1D3* cluster length and that passed QC are compared for Clusters 1 and 2. **B.** Potential disruptive human inversion haplotypes. The sequence structure of five haplotypes from two phylogenetically distinct clades (Clade 1 and Clade 2) are compared (SVbyEye) to the major configuration observed in humans. The two inversion haplotypes disrupt the potential fusion transcript by inserting a TBC1D3 paralog copy in reverse orientation with respect to the *NPEPPSP1* promoter. **C.** Comparative *TBC1D3* expression. Matched Kinnex transcriptome data of *NPEPPSP1-TBC1D3* transcripts were aligned to both haplotypes, and alignment score differences were computed and illustrated for inversion carriers and directly configured haplotypes. Inversion heterozygous genotypes consistently show a skew in alignment toward the direct orientation haplotypes, which maintain *NPEPPSP1-TBC1D3* orientation. Analysis of a homozygous inversion lymphoblastoid cell line shows a significantly reduced overall level of expression.

During this analysis we identified 16 samples where the terminal *TBC1D3* copy was inverted with respect to the *NPEPPS1* promoter (Fig. 6B). This inversion haplotype was found exclusively among samples of African origin (Supplementary Table 6) with an estimated allele frequency of 3.7% (16/437). Phylogenetic and comparative sequence analysis among humans predicts that this 76 kbp inversion potentially occurred twice (Fig. 6B; Supplementary Fig. 16). Because the inversion would, in principle, disrupt expression of the fusion transcript by placing *TBC1D3* and *NPEPPSP1* promoter on opposite strands, we assessed matched Kinnex transcriptome data from lymphoblast cell lines for six heterozygotes with the major allele (directly oriented) and the inverted allele as compared to homozygotes with the directly orientated allele—which represents the majority of human samples (Methods). Our analysis shows a skew in allelic expression, with *TBC1D3* transcripts being ∼13-42 times more likely to be assigned to the directly orientated allele when compared to the inverted allele in four heterozygous inversion samples (HG03453, HG02886, HG02922, NA19468; Fig. 6C). Additionally, these skewed samples showed matched reduction in both total *TBC1D3* and *NPEPPSP1-TBC1D3* fusion expression, a reduction of total and fusion expression of 22.7% and 22.1%, respectively. However, two of the six genotypes did not show an observable reduction in *TBC1D3* when compared to four controls (Fig. 6C).

Notably, we identified a single individual (NA18879) who was homozygous for the inversion and, as predicted, exhibited the greatest reduction in both total *TBC1D3* and *NPEPPSP1-TBC1D3* fusion transcript expression among the 11 samples analyzed. However, the homozygous inversion did not completely abolish fusion transcript expression, as we detected four *NPEPPSP1-TBC1D3* fusion transcripts in this individual (Figure 6C). Although the inversion is associated with an approximately 32 kbp insertion, the downstream *NPEPPSP1* and *TBC1D3* sequences retain their relative orientation with respect to the *NPEPPSP1* promoter. We hypothesize that this insertion reduces transcriptional efficiency but is ultimately removed through splicing, resulting in reduced expression rather than completely abolished *NPEPPSP1-TBC1D3* expression. Further functional experiments are needed to test this hypothesis. To investigate the prevalence and potential phenotypic consequences of this inversion, we identified eight tagging single-nucleotide polymorphisms in the unique sequence flanking the inversion, all in complete linkage disequilibrium with the inversion polymorphism (Supplementary Table 7). Using these markers, we identified 89 UK Biobank participants predicted to be homozygous for the inversion, predominantly among individuals who self-identified as African (69/89; Supplementary Table 8). Examination of their associated electronic health records revealed no obvious or consistent phenotype associated with inversion homozygosity.

## DISCUSSION

This work advances our understanding of *TBC1D3* regulation and evolution in four major ways. First, we demonstrate that *TBC1D3* expression in humans is regulated via a position effect, primarily as a result of a fusion duplication event with the upstream duplicated gene, *NPEPPSP1*. Paralog-specific regulation via position effect is a well-documented phenomenon, as exemplified in the locus control region (LCR) regulating immediately proximal green and red opsin paralogs in chromosome X, and none of the additional copies downstream (Hayashi et al., 1999). Other examples include the beta-globin LCR (Reik et al., 1998) and the 5’ LCR in *hGh* (Ho et al., 2006). Analogously, *NPEPPS1* defines an LCR of the terminal paralog of *TBC1D3* Cluster 2 with other paralogs being epigenetically repressed (Figs. 1 and 2).

Second, we identify the origin of this regulatory fusion as an SD event that duplicated ∼120 kbp—encompassing the *NPEPPS* promoter and first 11 exons. The ancestral *NPEPPS* locus maps 9.16 Mbp proximally to the *TBC1D3* Cluster 2 locus on chromosome 17 (Fig. 3). Gene fusion arising from the juxtaposition of partial SDs has been proposed as a relatively common mechanism of neofunctionalization, particularly in primates (Marques-Bonet, Girirajan et al., 2009; Marques-Bonet, Kidd, et al., 2009; Yoo et al., 2025). For example, *HYDIN2* arose from partial duplication of *HYDIN*, resulting in a novel gene fused to *LOC101927468* (Dougherty et al., 2017) and *CHRFAM7A* (i.e., *dupα7*), a human-specific fusion *CHRNA7* exons 5-10, and *FAM7A* exons A-E, regulating *CHR7A* in a dominant negative fashion (Sinkus et al., 2016).

In humans, the terminal copy of *TBC1D3* in Cluster 2 co-opted the regulatory program of *NPEPPS* to drive transcription of a fusion transcript. *NPEPPS*, also known as puromycin-sensitive aminopeptidase (PSA), is a cytoplasmic aminopeptidase conserved across metazoans that is broadly expressed across tissues and plays a key role in protein metabolism and cell cycle regulation (McLellan et al., 1988). In the brain, the NPEPPS protein is enriched in the cerebellum and hippocampus, where it provides a protective effect against tauopathies and associated neurodegenerative diseases, including Parkinson’s and dementia (Karsten et al., 2006; Muraoka et al., 2020). Disease associations remain difficult for *TBC1D3*, owing to the high sequence identity between paralogs and the significant degree of structural heterozygosity among individuals (Guitart et al., 2024). However, we find that *NPEPPSP1-TBC1D3* expression is enriched in the brain, and in particular, increases in expression in the cerebellum as the brain develops into adulthood (Supplementary Fig. 17). *In vitro* experiments of *TBC1D3* propose numerous mechanisms of *TBC1D3* function, all of which increase cell proliferation (Frittoli et al., 2008; Wainszelbaum et al., 2008, 2012; Ju et al., 2016; Hou et al., 2021). The human cerebellum is unique among other brain regions for its delayed maturation and high degree of neuronal density, which holds ∼80% of brain neurons despite constituting only 10% of volume (Van Essen et al., 2018). Recent *in vitro* studies suggest *TBC1D3* both promotes neuronal proliferation and protracts synaptic plasticity and contributes to neoteny (Hou et al., 2021; Dong et al., 2024). We suggest *TBC1D3* may contribute to human cerebellar development through this novel transcriptional context or the human-specific carboxy-terminus (Guitart et al., 2024).

Gene fusions driven by transcriptional readthrough often join two reading frames into a single novel protein (McCartney et al., 2019). In contrast, *NPEPPSP1-TBC1D3* retains two large, independent ORFs within a single transcript (Supplementary Fig. 9A). Translation of downstream ORFs is uncommon but does occur in eukaryotes through mechanisms including ribosomal reinitiation and cap-independent ribosome recruitment, although the unusually long 479-aa *NPEPPSP1* ORF may make efficient reinitiation less likely (Vattem et al., 2004; Wethmar et al., 2010; Gunišová et al., 2018; Haizel et al., 2020). Importantly, Ma et al. (2026) independently observed regularly distributed Ribo-seq footprints across the downstream *TBC1D3* ORF and detected TBC1D3 protein following transfection of *NPEPPS–TBC1D3* fusion cDNA constructs into HEK293T cells. These findings suggest downstream translation of *TBC1D3* does occur, though its translation is modest compared to *TBC1D3*-only controls. Low abundance of regulatory proteins, such as TBC1D3, however, can still substantially affect a cellular state, suggesting that limited TBC1D3 production from these transcripts could remain biologically meaningful (Gillespie et al., 2020).

Third, we identify developmentally regulated changes in *TBC1D3* isoform usage during neuronal differentiation. In our differentiation system, both *NPEPPSP1–TBC1D3* fusion transcripts and, more prominently, *TBC1D3*-solo transcripts increase as human stem cells differentiate into neurons, whereas *NPEPPSP1*-solo expression remains relatively stable (Fig. 5; Supplementary Figs. 11–13). The increased production of the *NPEPPSP1–TBC1D3* fusion transcript during neuronal differentiation may reflect developmentally regulated competition between polyadenylation and cleavage at the novel *NPEPPSP1* poly(A) site and continued transcription and splicing into *TBC1D3*. Determining the relevant cis-regulatory elements and trans-acting factors will require direct functional studies.

As these cells differentiate toward a neuronal fate, we wondered if these *TBC1D3*-solo transcripts truly derived from the same terminal *TBC1D3* genomic paralog. We initially considered whether these transcripts originated from nonterminal *TBC1D3* paralogs. However, matched transcriptome-to-assembly mapping and epigenetic profiling across multiple cell types indicate that all other *TBC1D3* paralogs are minimally expressed and remain epigenetically repressed (Figs. 1 and 2; Supplementary Table 6). Thus, the developmental increase in *TBC1D3*-solo expression is most consistent with transcription arising from the terminal *TBC1D3* locus, although the mechanism generating transcripts lacking the upstream *NPEPPSP1* sequence remains unresolved.

One possibility is that longer *NPEPPSP1–TBC1D3* transcripts undergo posttranscriptional processing that generates a *TBC1D3*-solo RNA. Such processing would remove the original 5’ cap and potentially render the transcript susceptible to RNA degradation. We tested whether *TBC1D3*-solo transcripts were preferentially degraded using cycloheximide treatment, disrupting translation-dependent RNA decay (Shoemaker & Green, 2012). Instead, cycloheximide preferentially increased *NPEPPSP1*-solo and *NPEPPSP1–TBC1D3* fusion transcripts relative to *TBC1D3*-solo transcripts, arguing against preferential degradation of the *TBC1D3*-solo isoforms (Supplementary Fig. 10). Alternatively, these *TBC1D3*-solo transcripts could arise through an internal or cryptic promoter within the terminal locus or through another, presently unresolved, RNA-processing mechanism. Determining how these *TBC1D3*-solo transcripts are generated, and whether their distinct 5’ architecture affects translation efficiency, stability, or subcellular localization, will require additional transcript- and protein-level investigation.

Finally, we place this regulatory architecture in an ape evolutionary framework. The *NPEPPSP1* duplication event occurred between 8–9 MYA, in the common ancestor of African apes. This time interval corresponds to a period of rapid chromosomal restructuring, including large-scale inversions and fusions that dramatically restructured local chromosomal regions in different ape lineages potentially as a result of extensive incomplete lineage sorting (Yang et al., 2025; Mao et al., 2021, Yoo et al., 2025). The *NPEPPSP1-TBC1D3* locus exhibits structural divergence across apes, with incomplete lineage sorting and lineage-specific rearrangements contributing to distinct haplotypes in humans, chimpanzees, and gorillas, including multiple large-scale and smaller inversion events (Supplementary Figs. 2, 3). Despite the shared origin of the fusion in all African great ape lineages, *TBC1D3* expression driven by *NPEPPSP1* appears to be human-specific. In the genus *Pan*, a subsequent 44 kbp chimpanzee duplication inserted *CCL4L2* between *NPEPPSP1* and *TBC1D3*, disrupting the fusion in both chimpanzees and bonobos. In gorilla, a 51 kbp deletion removed the *NPEPPSP1* promoter and initial exons, ablating the fusion transcript. Based on the genomic architecture, only humans are, thus, capable of producing the *NPEPPSP1-TBC1D3* fusion product at high levels during neurodevelopment.

We find that the expressed *TBC1D3* gene has been dramatically restructured during human evolution. Our previous work identified a human-specific C-terminal truncation resulting in a 58-aa modification in all expressed human copies. Here, we show that the 5’ end of the major human *TBC1D3* transcript is also unique to our lineage—it includes the first 11 exons of *NPEPPSP1*. Further, we provide evidence that this modification boosts expression during neuronal development. The functional consequences of these alterations—for translation, localization, or function—remain unresolved. Experimental perturbation of *TBC1D3* remains challenging due to its high sequence identity across paralogs. However, our findings present a potential solution: targeting the fixed *NPEPPSP1* promoter, which regulates the polymorphic *TBC1D3* cluster through a position effect. This regulatory mechanism—where a fixed regulatory element controls expression of structurally variable loci—may be more common than expected, offering a potential target for perturbation studies to better understand the regulation and importance of these structurally variable and medically relevant genes that have, until recently, been impossible to characterize.

## SUPPLEMENTARY FIGURES

**Supplementary Figure 1.**
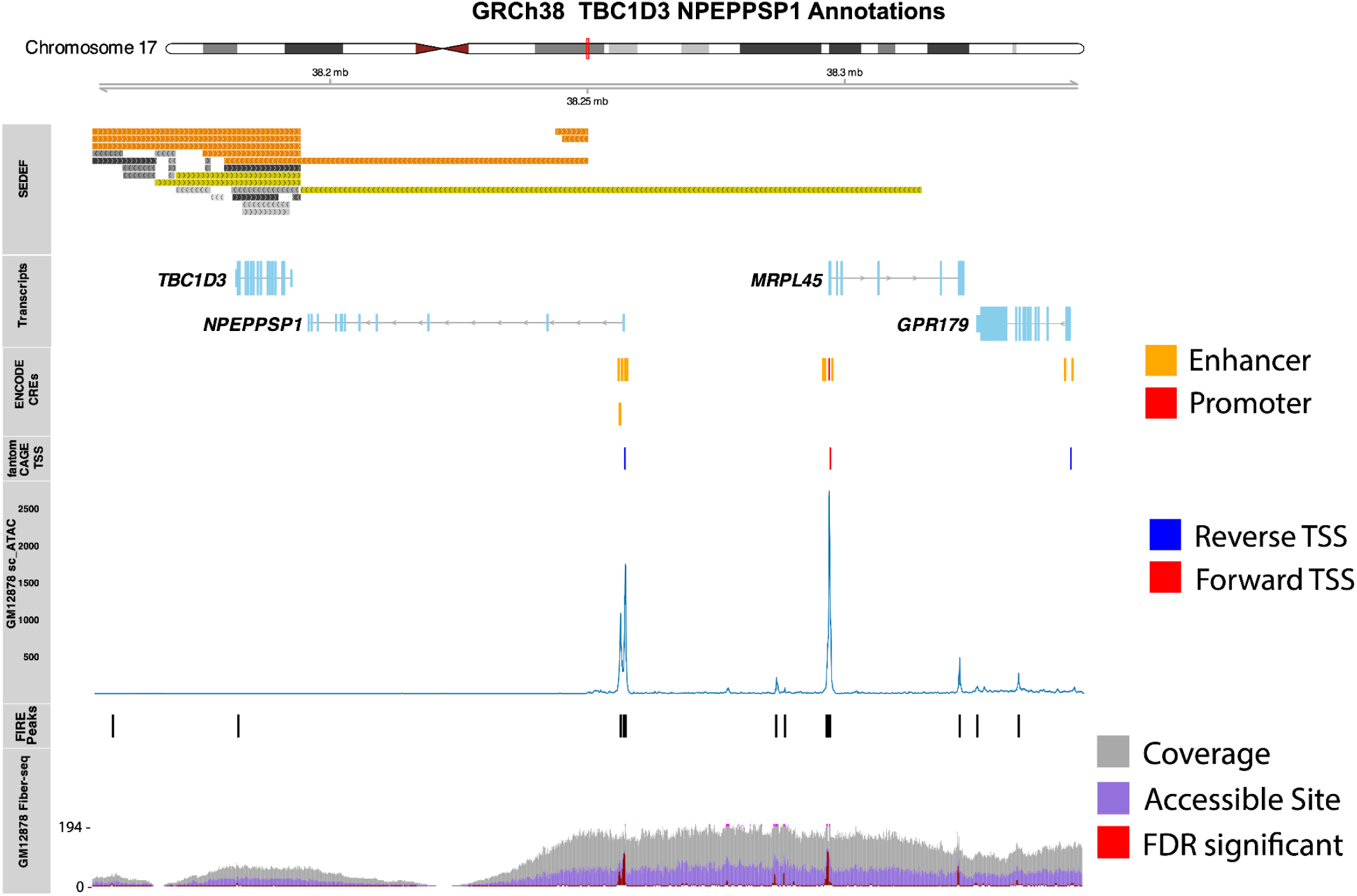
Gene regulatory elements in GRCh38. Regulatory elements identified by the ENCODE project and scATAC are compared to Fiber-seq data of the GM2878 cell line (Vollger et al., 2025). TSS: transcription start site; CRE: cis-regulatory element.

**Supplementary Figure 2.**
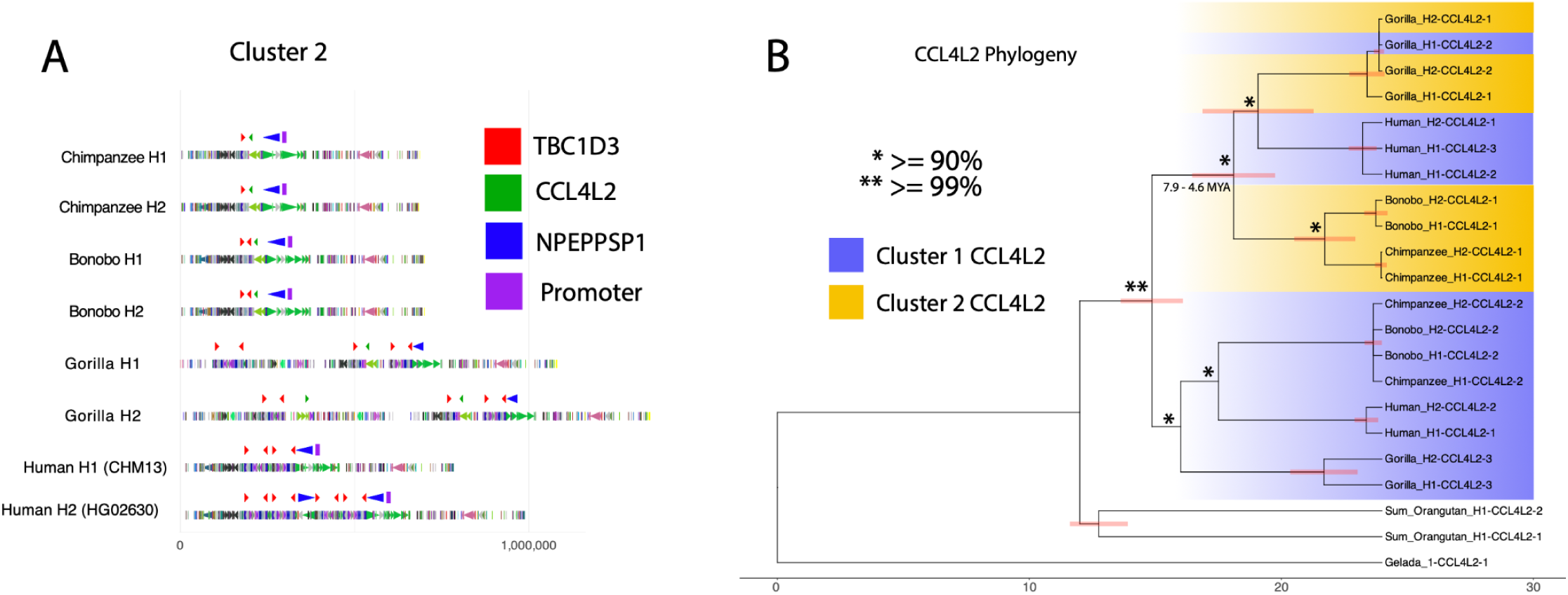
*CCL4L2* structure and phylogeny. **A.** Genomic architecture of *TBC1D3* Cluster 2 in African apes. *CCL4L2* obstructs *NPEPPSP1* from *TBC1D3* and co-opts fusion expression in the *Pan* genera. This structure is fixed in both chimpanzee and bonobo. **B.** Maximum likelihood phylogeny of *CCL4L2* sequence. Timing estimates of the *CCl4L2* obstruction event are between 4.6 and 7.9 MYA, and human and gorilla *CCL4L2* copies are more similar to one another than *Pan*.

**Supplementary Figure 3.**
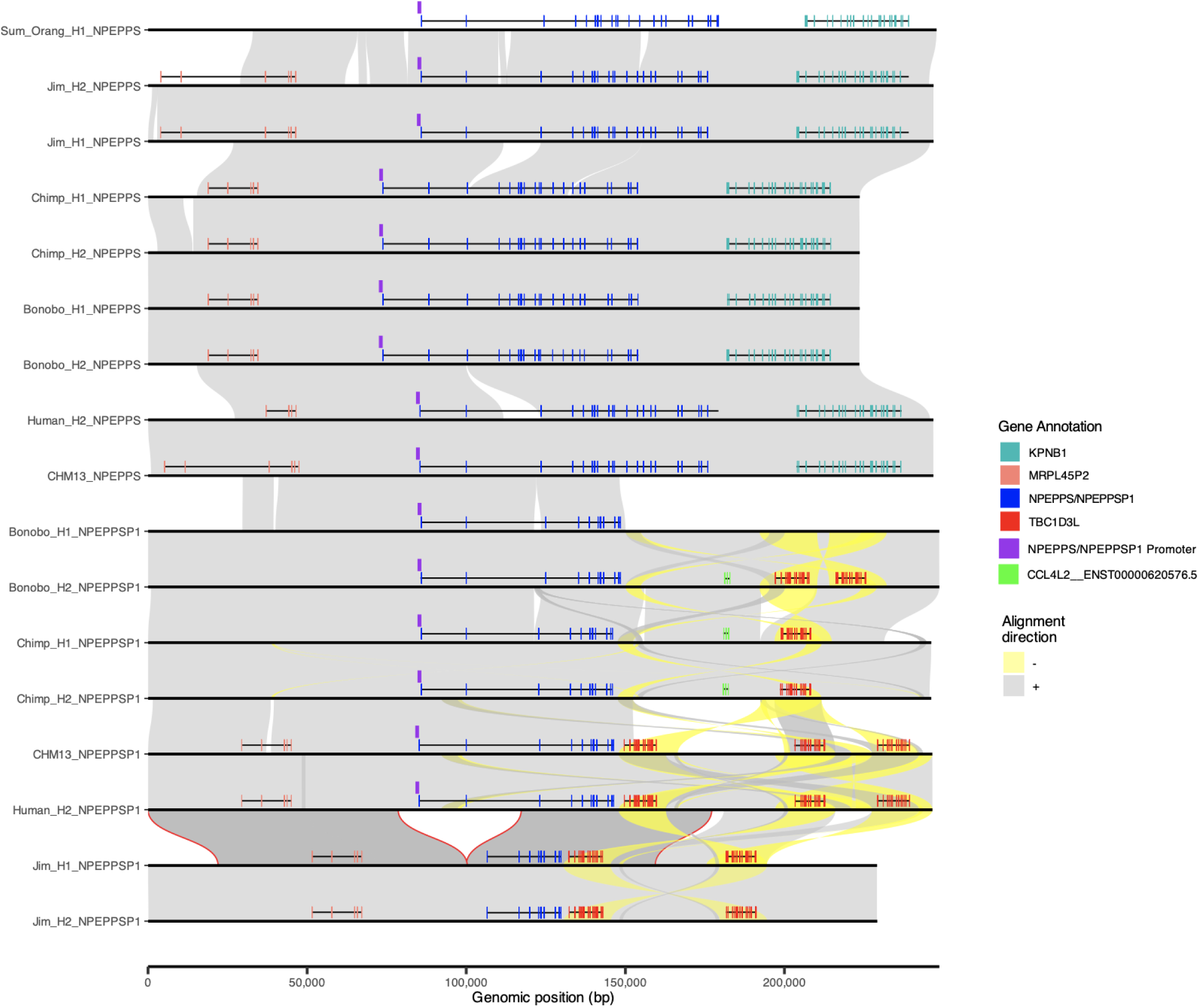
Gorilla deletion. Alignments of the *NPEPPS* (ancestral) and *NPEPPSP1-TBC1D3* (derived) loci are illustrated, showing the orthologous *NPEPPSP1* promoter inherited across African apes but lost in the gorilla lineage.

**Supplementary Figure 4.**
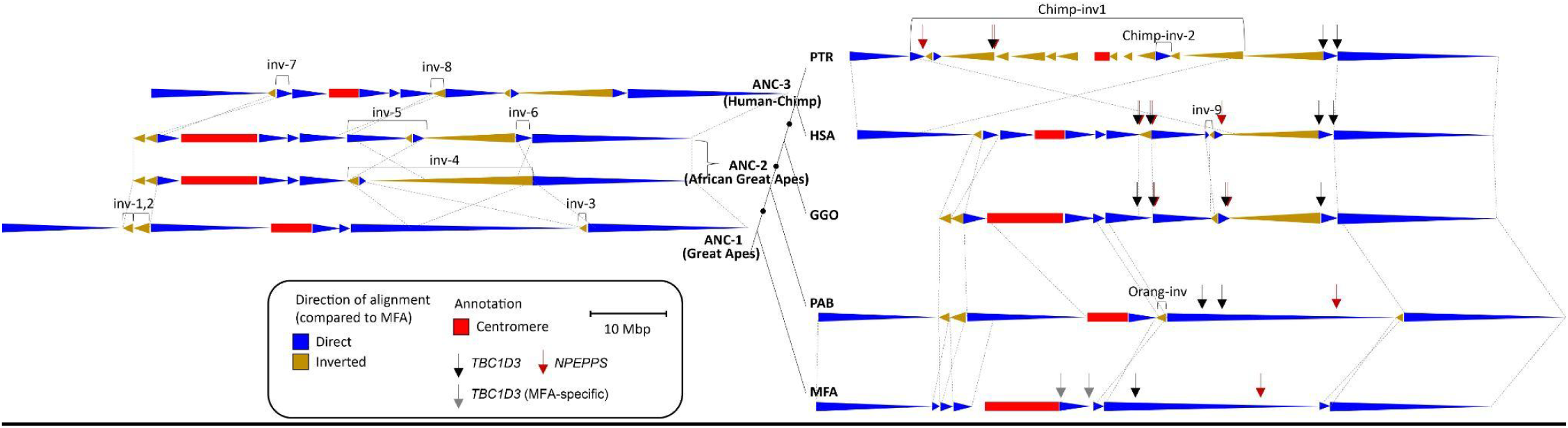
Comparative analysis of primate inversions. The location of 12 inversions, the breakpoints, and their evolutionary origin based on comparative sequence analysis of six primate genomes (left) projected onto a generally accepted phylogeny (right). Syntenic (blue) and inverted (yellow) segments and centromere locations (red) with respect to macaque. The location of *TBC1D3* (black) and *NPEPPS* (burgundy arrows) duplicate genes with respect to breakpoints is shown. Inversions are labelled at the most ancestral position.

**Supplementary Figure 5.**
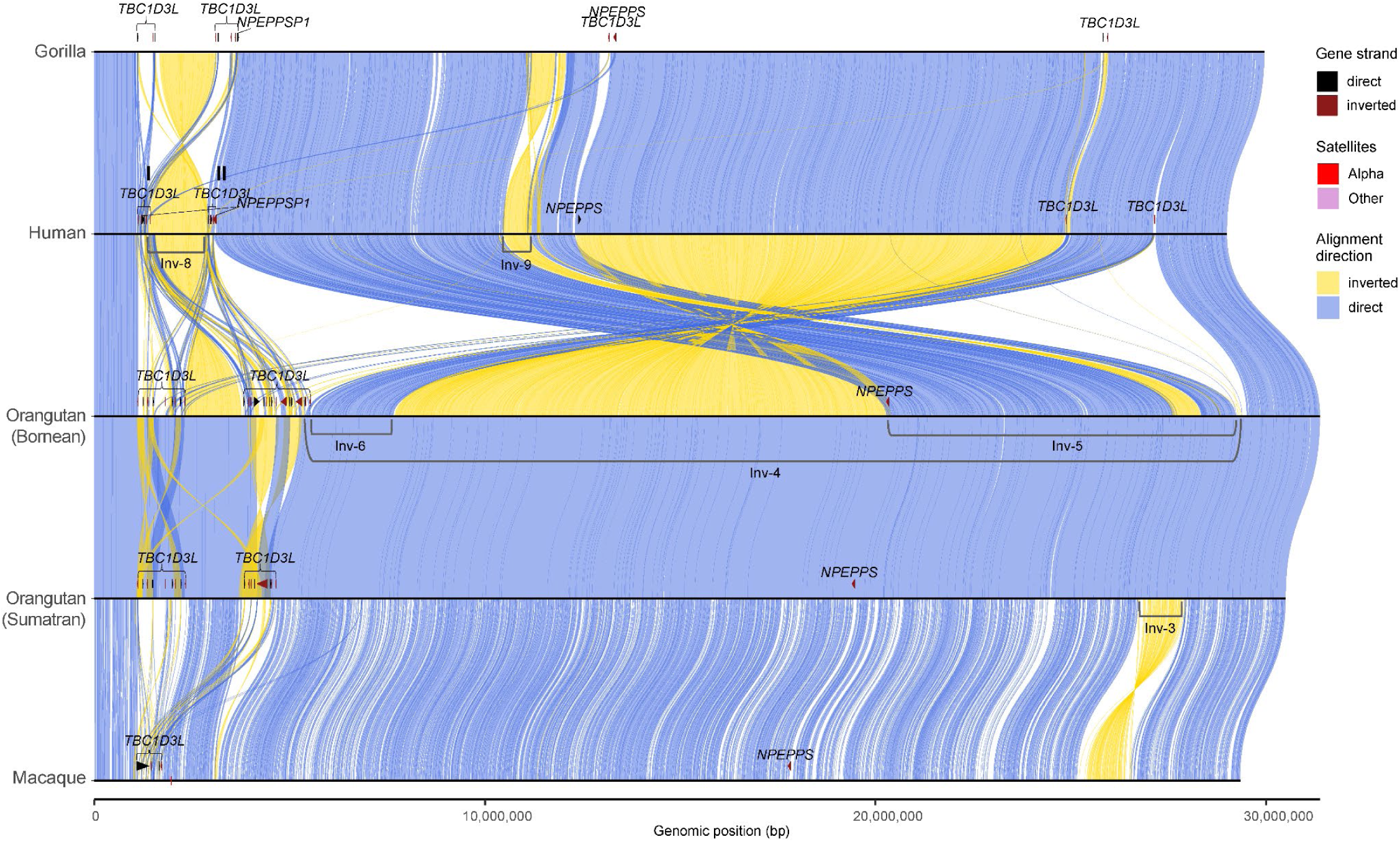
Alignment of zoomed-in *TBC1D3* locus (chr17:36-65Mbp) across primate lineage. The direct syntenic alignments are indicated by blue while the reverse complement alignments are indicated by yellow representing inversions. *TBC1D3* and *NPEPPSP1* are indicated above each alignment track colored by their strand, + in black and - in red. For human, *TBC1D3* Clusters 1 and 2 are indicated above the gene track. The inversions are annotated below each track.

**Supplementary Figure 6.**
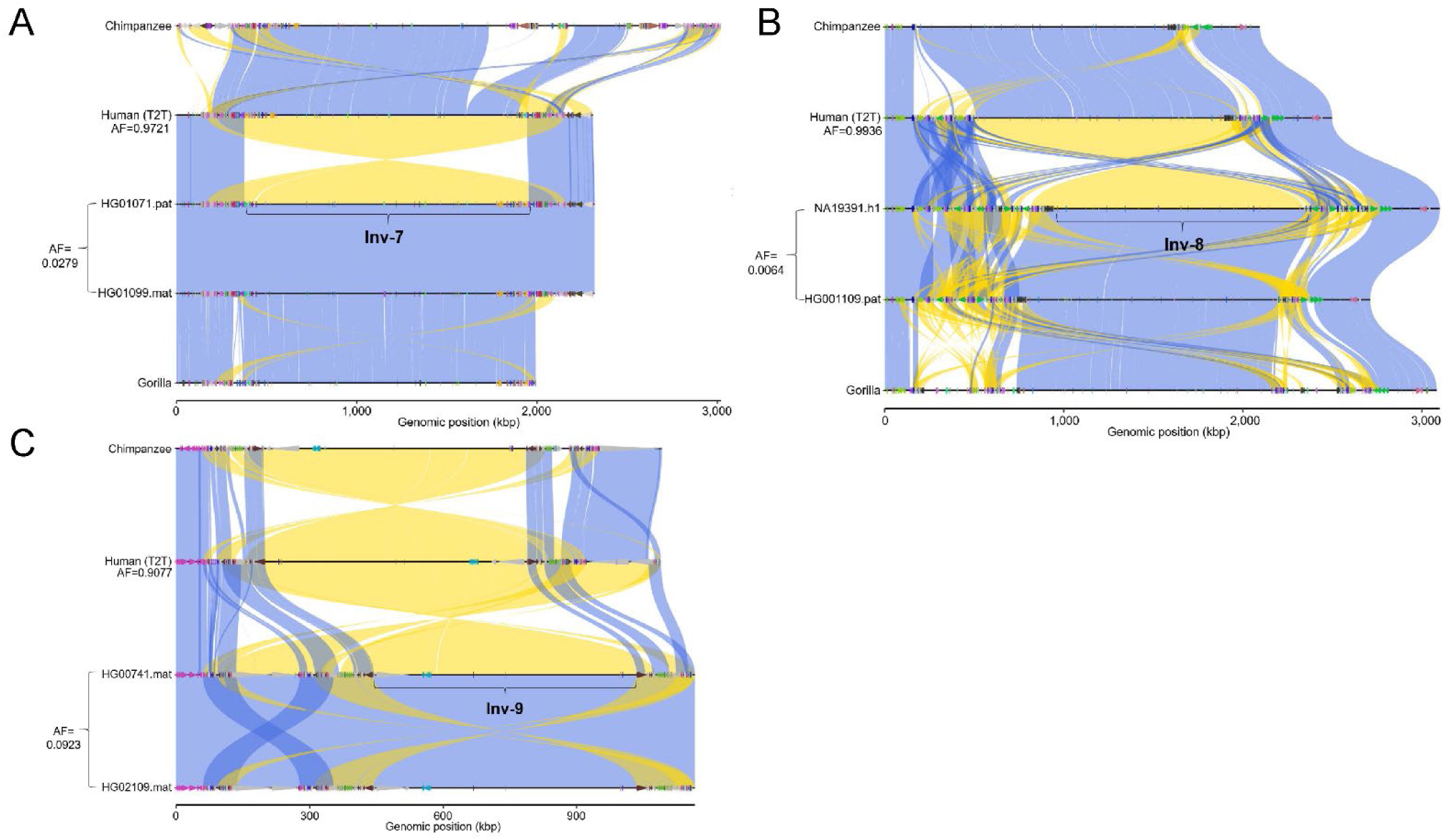
Human haplotypes with ancestral states. Inversions including **A)** inv-7, **B)** inv-8, and **C)** inv-9, which contain allele frequency among the human population are displayed. Two representative haplotypes of human populations are shown for each inversion, with allele frequencies indicated on the left. Duplicons predicted by DupMasker are shown at the center illustrating the duplication structure that is flanking each inversion.

**Supplementary Figure 7.**
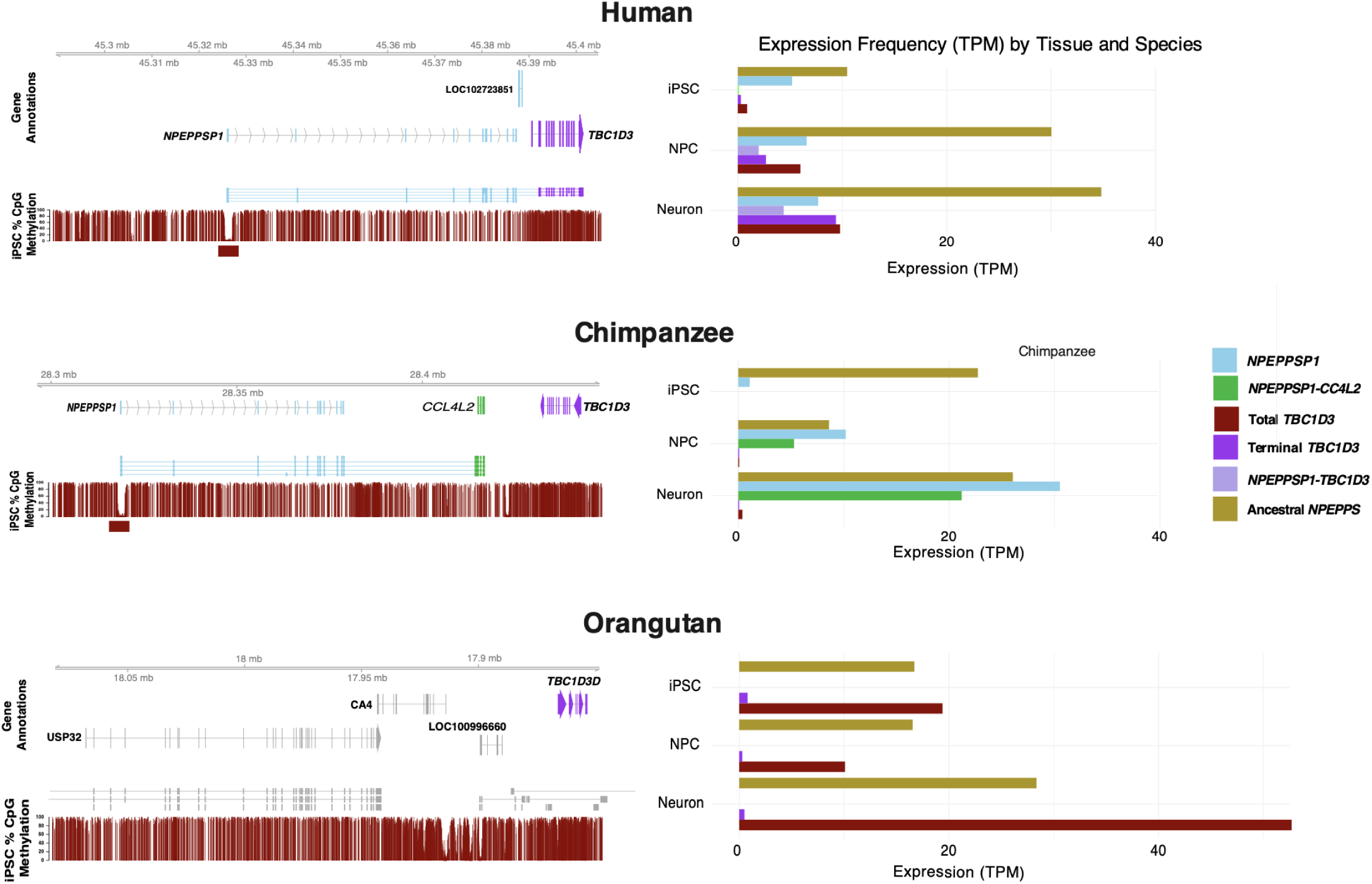
Comparative expression with *NPEPPS* and total *TBC1D3* in a neuronal developmental cell culture model of great apes. Gene annotation and methylation of *NPEPPSP1-TBC1D3* can be observed on the left for human (top), chimpanzee (middle), and orangutan (bottom). On the right, expression of *NPEPPSP1*, terminal *TBC1D3*, and their fusion are compared to global *TBC1D3* and *NPEPPS* expression, normalized as transcripts per million (TPM; Methods).

**Supplementary Figure 8.**
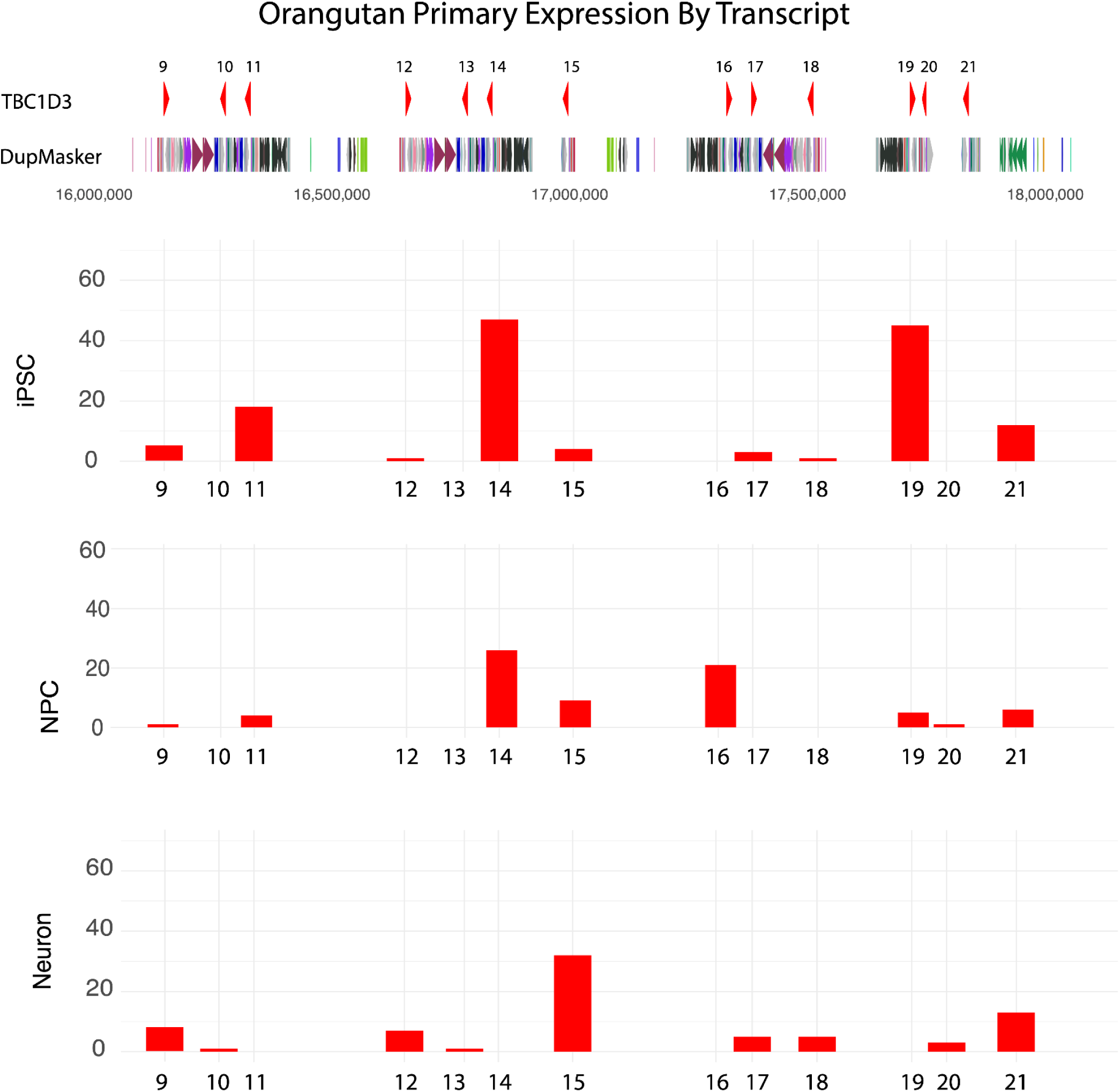
Orangutan expression by *TBC1D3* transcript. Orangutan primary transcripts map to numerous internal *TBC1D3* paralog copies (*TBC1D3*-14,15,16,19).

**Supplementary Figure 9.**
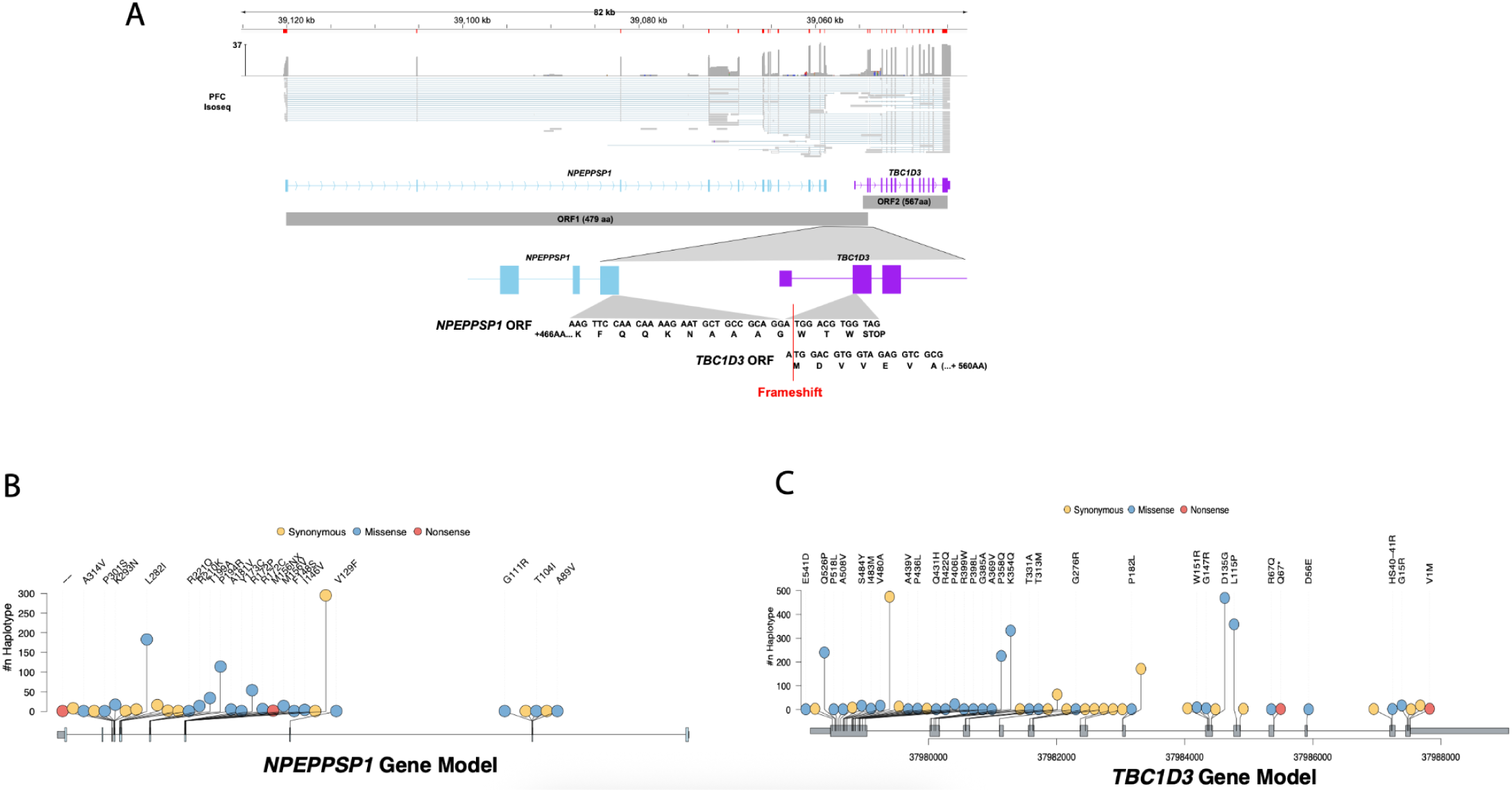
*NPEPPSP1-TBC1D3* retains two ORFs. **A.** Iso-Seq data from the prefrontal cortex illustrates the fusion transcript across *NPEPPSP1* and *TBC1D3*. Below are two open reading frames (ORFs) identified with ORFfinder (Methods), which span *NPEPPSP1* and *TBC1D3* in separate reading frames overlapping by 13 nucleotides. **B.** Lollipop plots of *NPEPPSP1* illustrate synonymous (yellow), missense (blue), and nonsense (red) variants. Three missense variants were identified in 2,013 genome assemblies. **C.** Same as (B) but with *TBC1D3* ORF2. One missense variant was identified across the gene body.

**Supplementary Figure 10.**
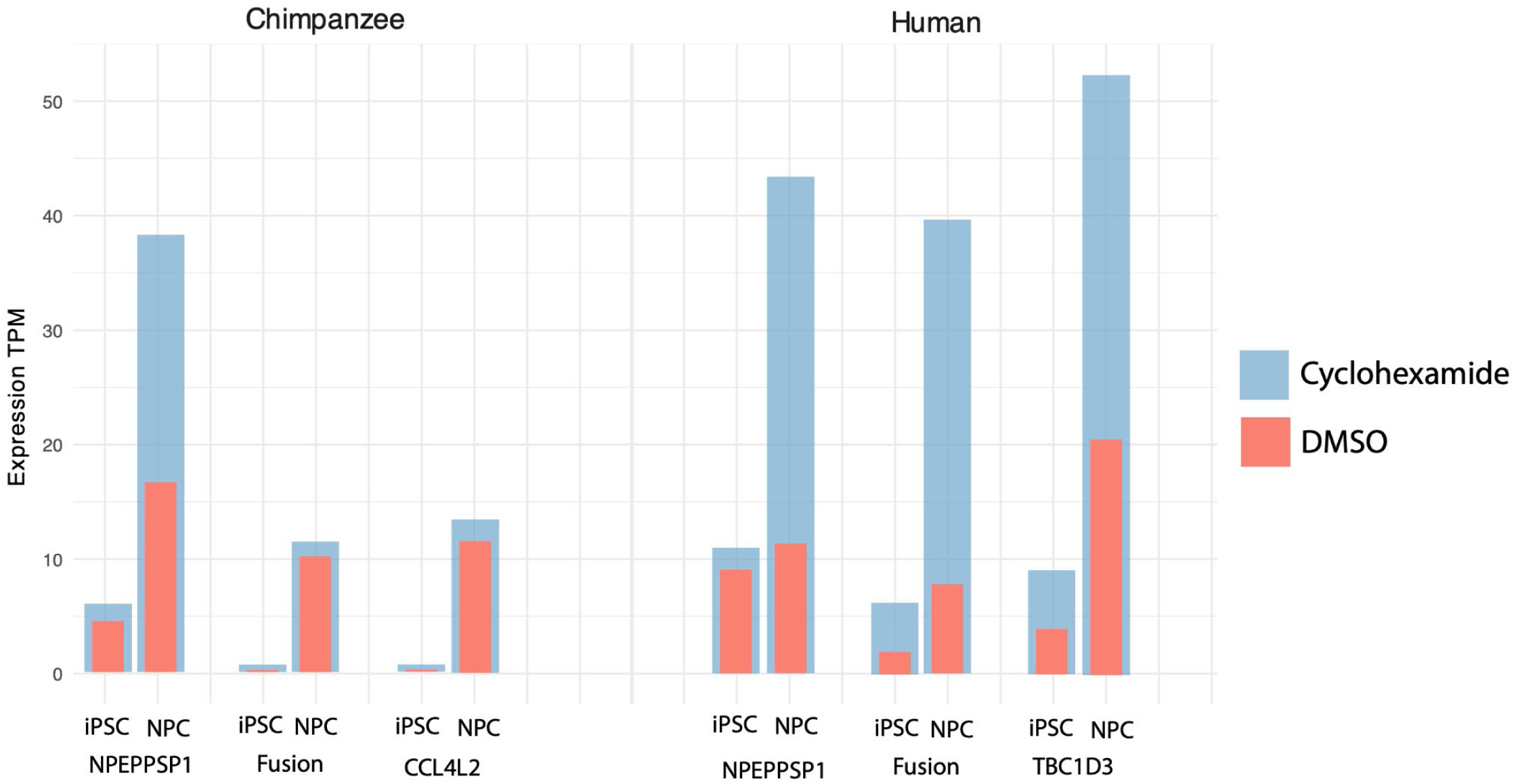
Disruption of translation-dependent degradation. iPSCs and NPCs from chimpanzee and humans were treated with either DMSO or cycloheximide, a disruptor of translation-dependent degradation, to investigate posttranscriptional fate of fusion genes. Notably, *NPEPPSP1* is preferentially rescued in both species in NPCs relative to *TBC1D3*, though in humans *NPEPPS1*, *TBC1D3*, and the fusion isoform all increase equally.

**Supplementary Figure 11.**
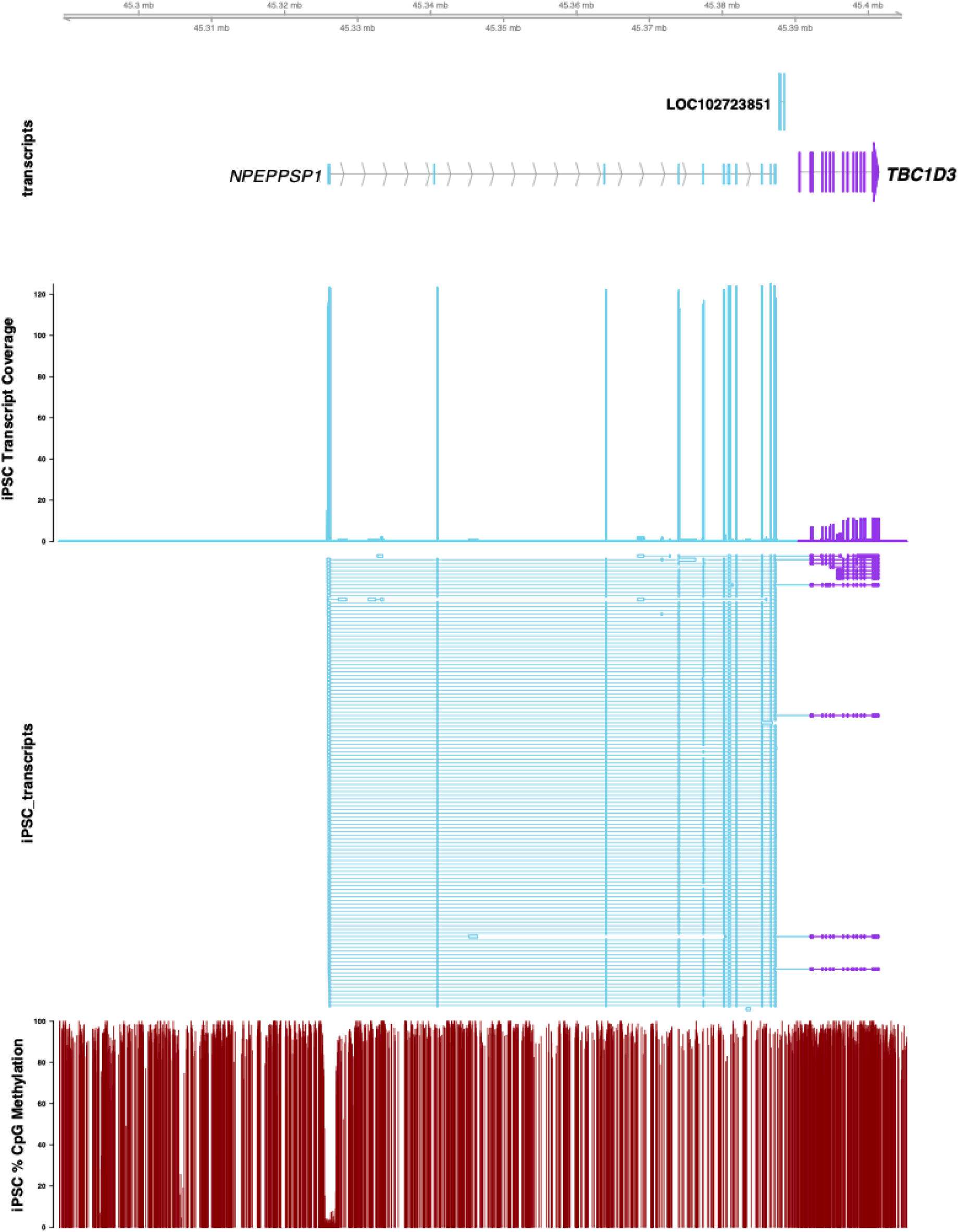
Human iPSC expression and methylation. Human iPSC gDNA and full-length transcripts were mapped back to their donor-specific genome assembly (DSA) and annotated for *NPEPPSP1* (blue) or *TBC1D3* (purple) gene models. The *NPEPPSP1* promoter may be seen as a dip in methylation at *NPEPPSP1* exon 1.

**Supplementary Figure 12.**
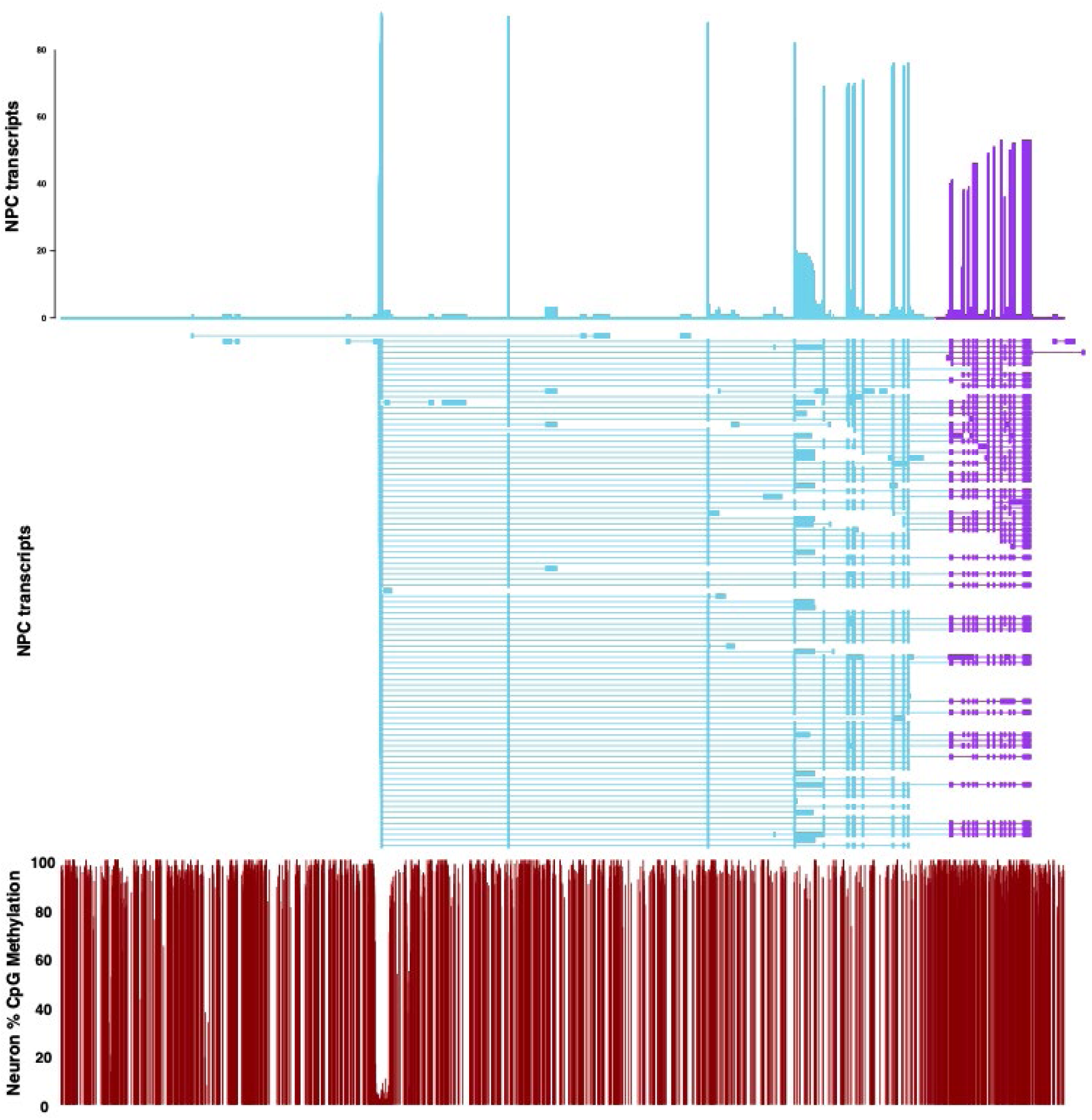
Human NPC expression and methylation. Human NPC gDNA and full-length transcripts were mapped back to their DSA and annotated for *NPEPPSP1* (blue) or *TBC1D3* (purple) gene models. The *NPEPPSP1* promoter may be seen as a dip in methylation at *NPEPPSP1* exon 1.

**Supplementary Figure 13.**
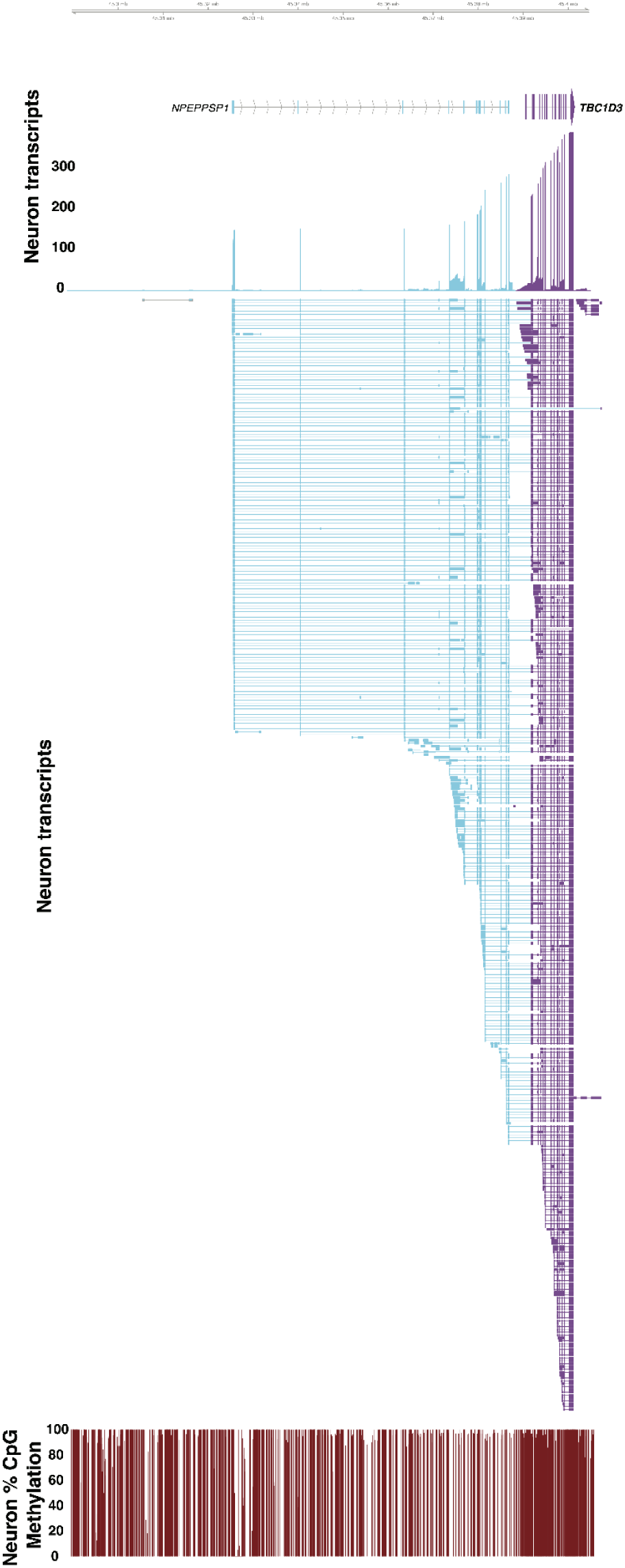
Human neuron expression and methylation. Human neuron gDNA and full-length transcripts were mapped back to their DSA and annotated for *NPEPPSP1* (blue) or *TBC1D3* (purple) gene models. The *NPEPPSP1* promoter may be seen as a dip in methylation at *NPEPPSP1* exon 1. Notably, in neurons, *TBC1D3* has increased in expression relative to *NPEPPSP1*, in contrast to NPCs or iPSCs.

**Supplementary Figure 14.**
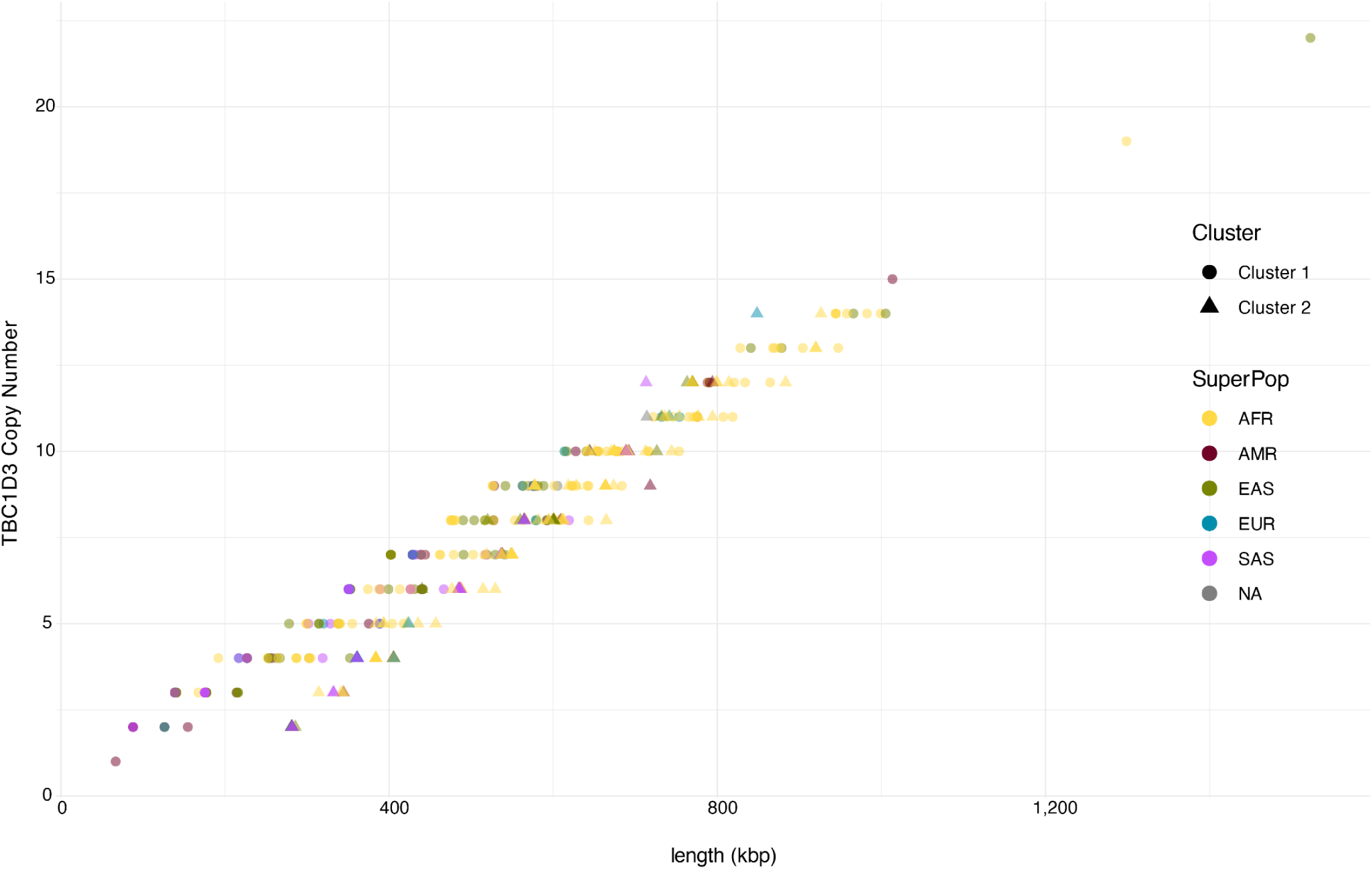
Cluster 1 and Cluster 2 size versus copy number. Clusters 1 and 2 show broadly similar distribution of size and *TBC1D3* copy number, though Cluster 1 extremes are more significant, and Cluster 1 has a slightly larger average (6.8 copies) relative to Cluster 2 (6.2 copies, *p* = 0.004, Welch’s *t*-test).

**Supplementary Figure 15.**
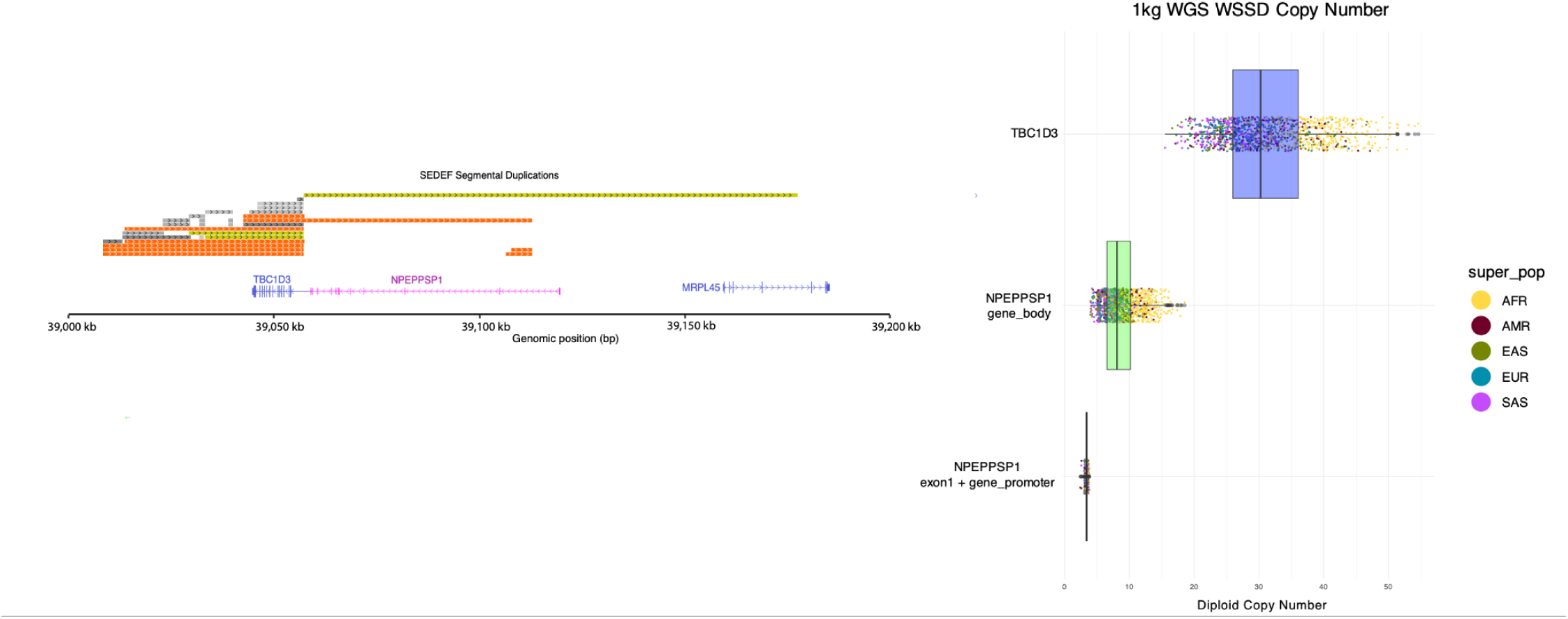
*NPEPPSP1* promoter fixed in copy number. Left: Segmental duplication (SD) annotation of the *NPEPPSP1-TBC1D3* locus illustrates the numerous high-identity SDs across both *TBC1D3* and *NPEPPSP1* but does not include the *NPEPPSP1* exon 1 or its upstream promoter. Right: Diploid copy number estimates of *TBC1D3*, *NPEPPSP1*, and *NPEPPSP1* promoter in the 1000 Genomes Project (1KGP) identified by whole-genome shotgun sequence detection (WSSD; Sudmant et al., 2010) illustrate the fixed copy number of the *NPEPPSP1-TBC1D3* promoter relative to either *NPEPPSP1* or *TBC1D3*.

**Supplementary Figure 16.**
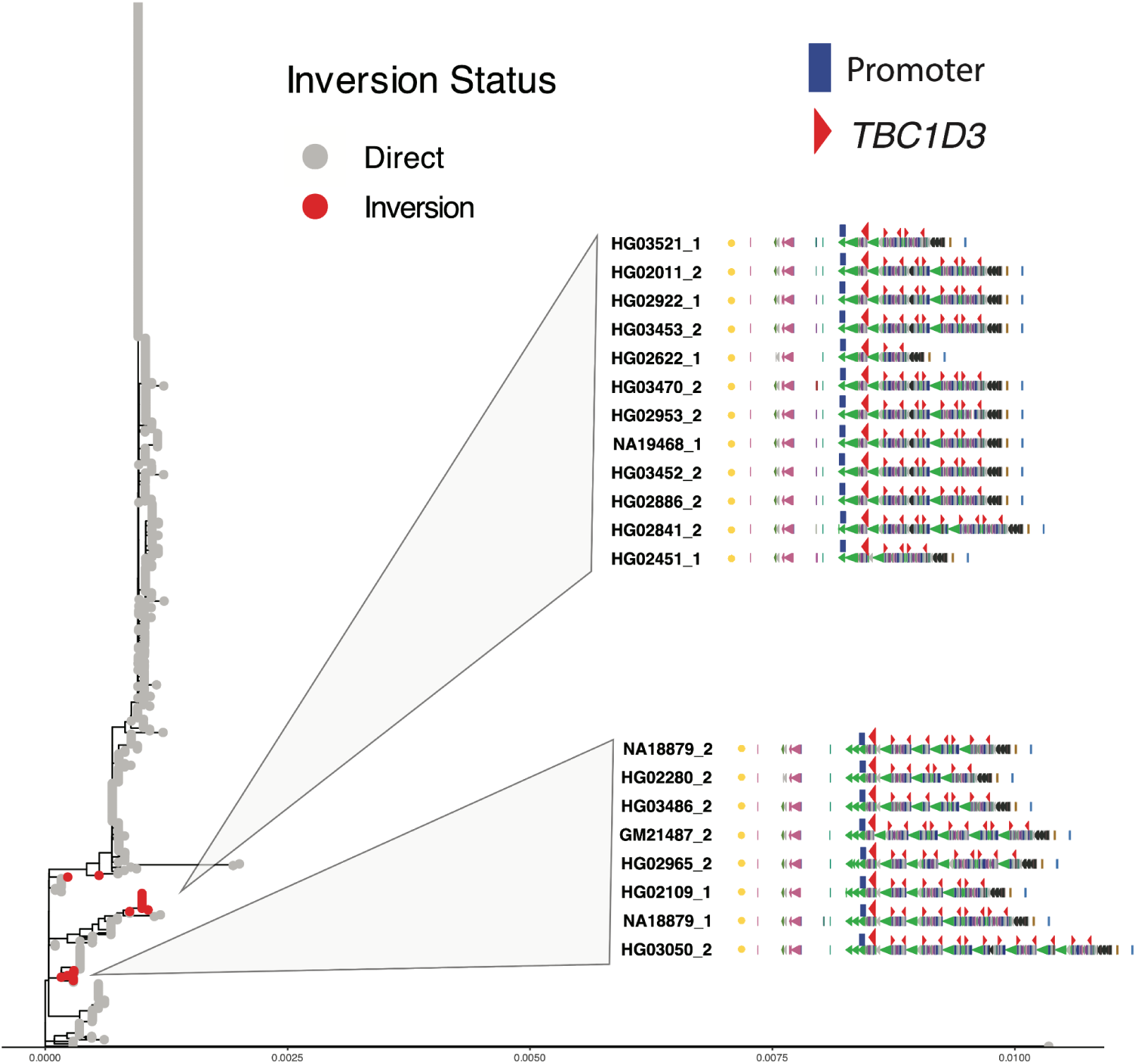
Two *TBC1D3* terminal inversion clades. A maximum likelihood sequence phylogeny of 20 kbp unique sequence directly upstream of *NPEPPSP1* for all assembled and validated assemblies shows haplotypes with terminal *TBC1D3* paralogs inverted relative to upstream *NPEPPSP1* promoter arose two separate times.

**Supplementary Figure 17.**
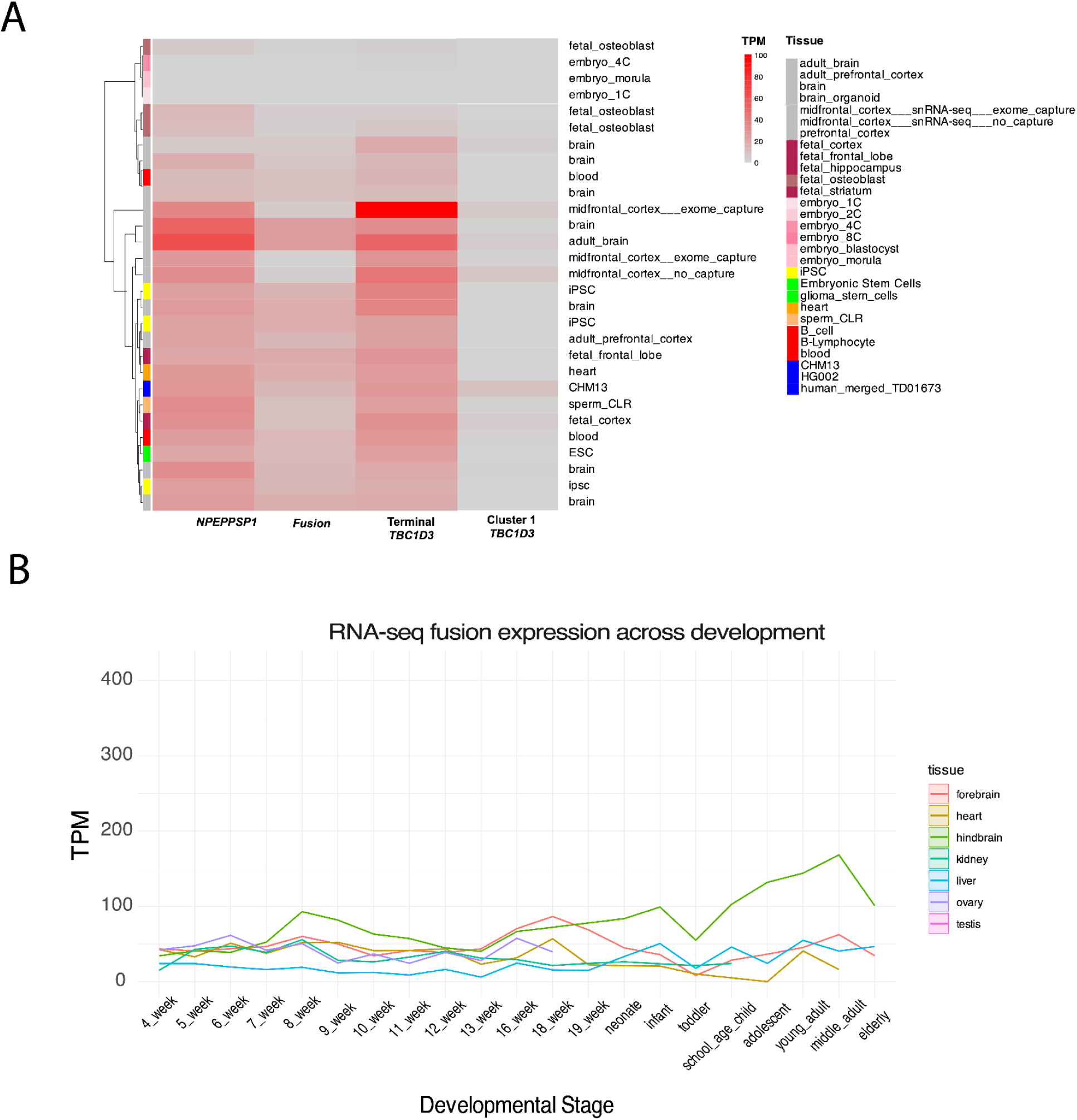
*TBC1D3* expression. **A.** Full-length transcriptome sequencing of various libraries mapped to *NPEPPSP1*, *NPEPPSP1-TBC1D3* fusion, terminal *TBC1D3*, or any Cluster 1 *TBC1D3* (control). **B.** RNA-seq of *NPEPPSP1-TBC1D3* fusion across development.

## SUPPLEMENTARY TABLES

**Supplementary Table 1.**
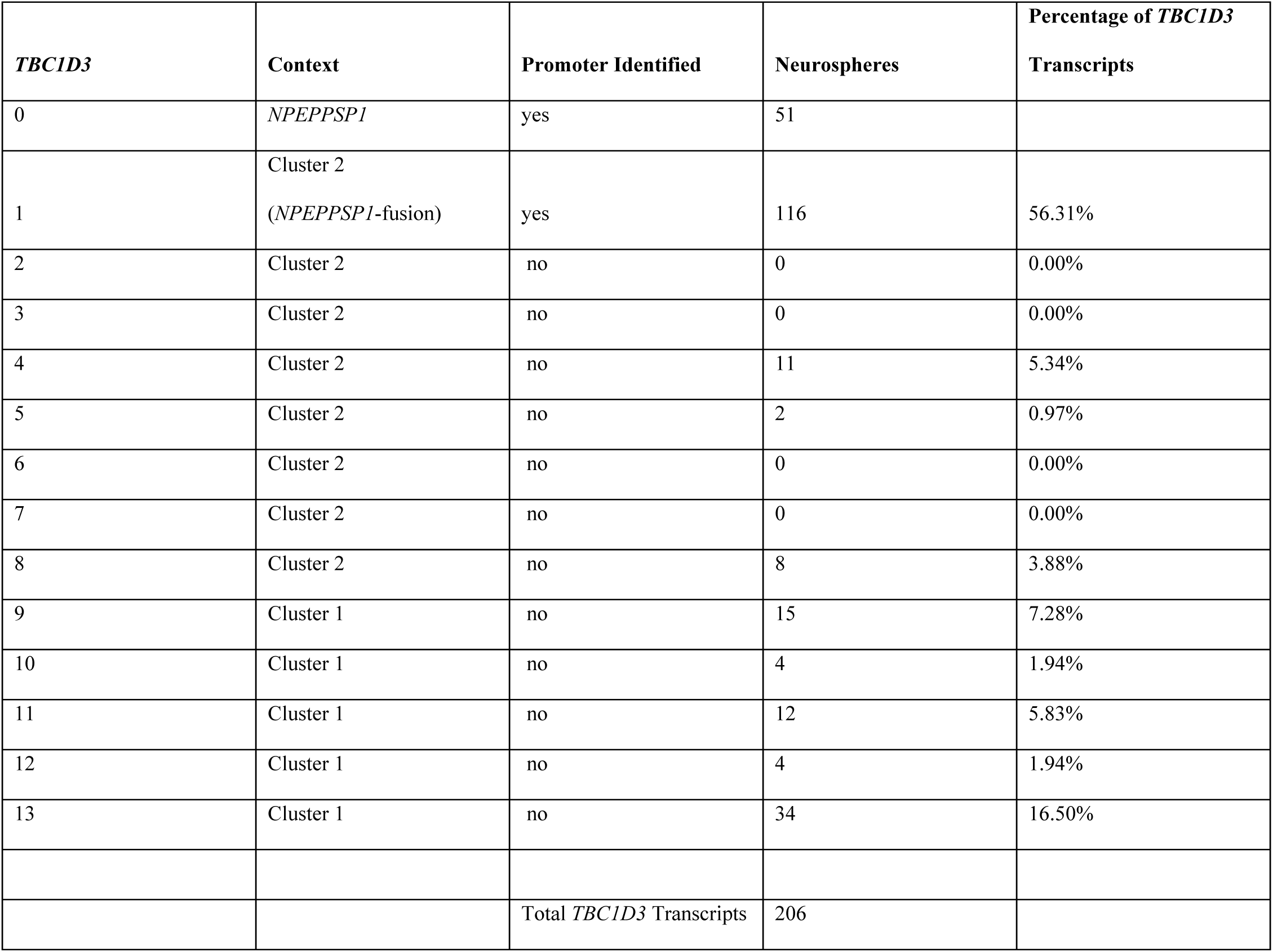
Neurospheres mappings by *TBC1D3* paralog.

**Supplementary Table 2.** Summary of inversions characterized in chr17.

| Inversion ID | Ancestral/HSA-derived | Lineage label | Control genome | Query genome | Query Chr | Query start | Query end | Size (bp) | Human allele count | Human allele freq (n=466) |
|---|---|---|---|---|---|---|---|---|---|---|
| Inv-1 | Ancestral | Great ape (ANC-1) | MFA | PAB | chr19_hap1_hsa17 | 16477588 | 17417479 | 939891 | 0 | 0.0000 |
| Inv-2 | Ancestral-recurrent | Great ape (ANC-1) | MFA | PAB | chr19_hap1_hsa17 | 17729085 | 19463215 | 1734130 | 13 | 0.0279 |
| Inv-3 | Ancestral | Great ape (ANC-1) | MFA | PAB | chr19_hap1_hsa17 | 75812956 | 77031830 | 1218874 | 0 | 0.0000 |
| Inv-4 | Ancestral | African Great ape (ANC-2) | PAB | GGO | chr4_mat_hsa17x5 | 30192433 | 54482959 | 24290526 | NA | NA |
| Inv-5 | Ancestral | African Great ape (ANC-2) | PAB | GGO | chr4_mat_hsa17x5 | 45109025 | 54482959 | 9373934 | 0 | 0.0000 |
| Inv-6 | Ancestral | African Great ape (ANC-2) | PAB | GGO | chr4_mat_hsa17x5 | 30192433 | 32271580 | 2079147 | 0 | 0.0000 |
| Inv-7 | Ancestral-recurrent | Human-Chimp ancestor (ANC-3) | GGO | HSA | chr17 | 16821891 | 18331721 | 1509830 | 13 | 0.0279 |
| Inv-8 | Ancestral | Human-Chimp ancestor (ANC-3) | GGO | HSA | chr17 | 37444979 | 38843266 | 1398287 | 3 | 0.0064 |
| Inv-9 | HSA-derived | Human (HSA) | GGO | HSA | chr17 | 46489489 | 47078697 | 589208 | 43 | 0.0923 |
| Chimp-inv-1 | PTR-derived | Chimpanzee (PTR) | HSA | PTR | chr19_hap1_hsa17 | 18542288 | 62681614 | 44139326 | NA | NA |
| Chimp-inv-2 | PTR-derived | Chimpanzee (PTR) | HSA | PTR | chr19_hap1_hsa17 | 50956137 | 52911392 | 1955255 | NA | NA |
| Orang-inv | PAB-derived | Orangutan (Sumatran; PAB) | MFA | PAB | chr19_hap1_hsa17 | 44783386 | 45744419 | 961033 | 0 | 0.0000 |

**Supplementary Table 3.**
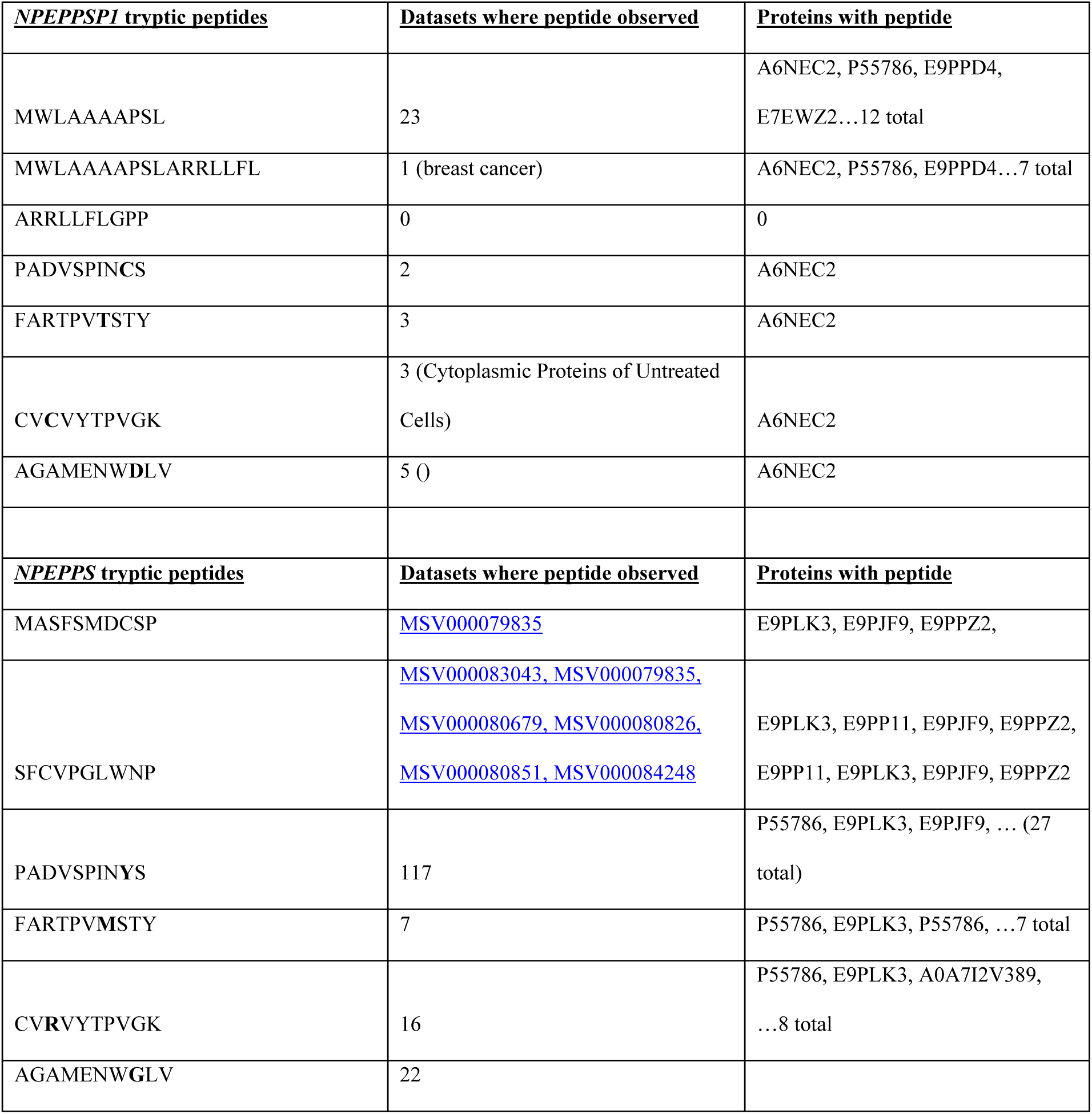
*NPEPPS* versus *NPEPPSP1* mass spectra peptide observations.

| ***NPEPPSP1*** **tryptic peptides** | **Datasets where peptide observed** | **Proteins with peptide** |
|---|---|---|
| MWLAAAAPSL | 23 | A6NEC2, P55786, E9PPD4, E7EWZ2…12 total |
| MWLAAAAPSLARRLLFL | 1 (breast cancer) | A6NEC2, P55786, E9PPD4…7 total |
| ARRLLFLGPP | 0 | 0 |
| PADVSPIN**C**S | 2 | A6NEC2 |
| FARTPV**T**STY | 3 | A6NEC2 |
| CV**C**VYTPVGK | 3 (Cytoplasmic Proteins of Untreated Cells) | A6NEC2 |
| AGAMENW**D**LV | 5 () | A6NEC2 |
| ***NPEPPS*** **tryptic peptides** | **Datasets where peptide observed** | **Proteins with peptide** |
| MASFSMDCSP | MSV000079835 | E9PLK3, E9PJF9, E9PPZ2, |
| SFCVPGLWNP | MSV000083043, MSV000079835, MSV000080679, MSV000080826, MSV000080851, MSV000084248 | E9PLK3, E9PP11, E9PJF9, E9PPZ2, E9PP11, E9PLK3, E9PJF9, E9PPZ2 |
| PADVSPIN**Y**S | 117 | P55786, E9PLK3, E9PJF9, … (27 total) |
| FARTPV**M**STY | 7 | P55786, E9PLK3, P55786, …7 total |
| CV**R**VYTPVGK | 16 | P55786, E9PLK3, A0A7I2V389, …8 total |
| AGAMENW**G**LV | 22 |  |

**Supplementary Table 4.**
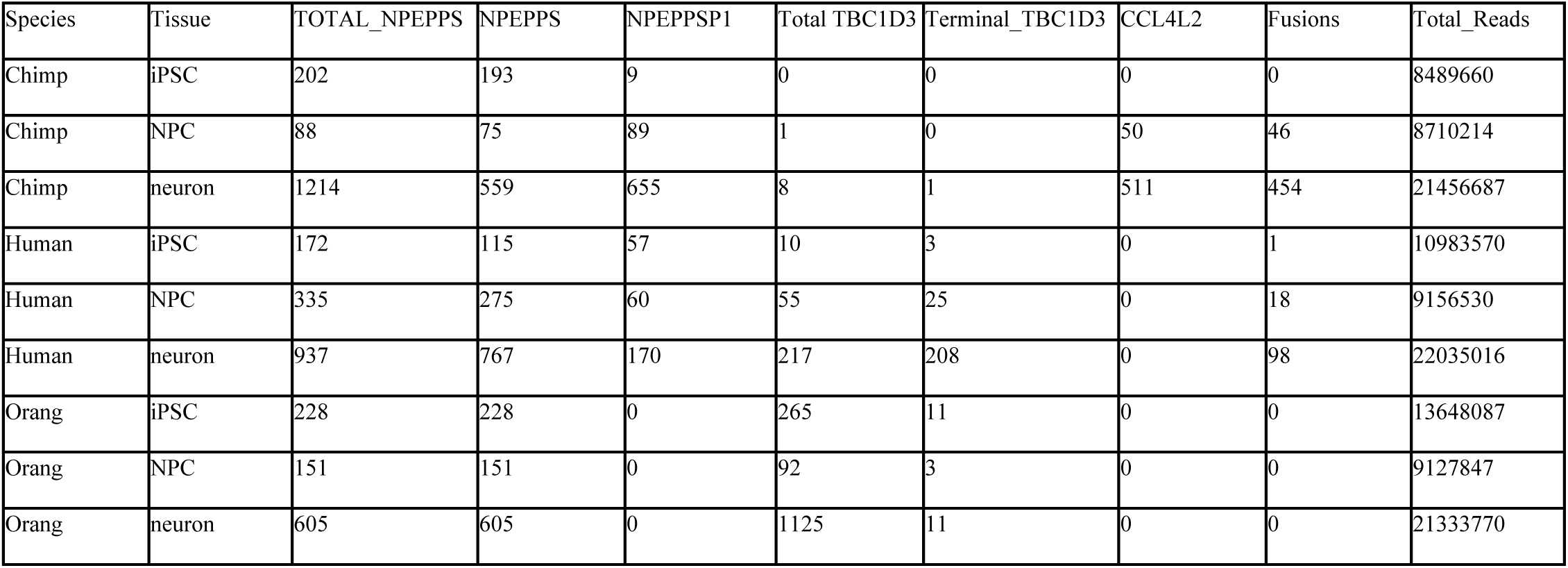
iPSC-NPC-iNeuron transcript counts.

| Species | Tissue | TOTAL_NPEPPS | NPEPPS | NPEPPSP1 | Total TBC1D3 | Terminal_TBC1D3 | CCL4L2 | Fusions | Total_Reads |
|---|---|---|---|---|---|---|---|---|---|
| Chimp | iPSC | 202 | 193 | 9 | 0 | 0 | 0 | 0 | 8489660 |
| Chimp | NPC | 88 | 75 | 89 | 1 | 0 | 50 | 46 | 8710214 |
| Chimp | neuron | 1214 | 559 | 655 | 8 | 1 | 511 | 454 | 21456687 |
| Human | iPSC | 172 | 115 | 57 | 10 | 3 | 0 | 1 | 10983570 |
| Human | NPC | 335 | 275 | 60 | 55 | 25 | 0 | 18 | 9156530 |
| Human | neuron | 937 | 767 | 170 | 217 | 208 | 0 | 98 | 22035016 |
| Orang | iPSC | 228 | 228 | 0 | 265 | 11 | 0 | 0 | 13648087 |
| Orang | NPC | 151 | 151 | 0 | 92 | 3 | 0 | 0 | 9127847 |
| Orang | neuron | 605 | 605 | 0 | 1125 | 11 | 0 | 0 | 21333770 |

**Supplementary Table 5.** iPSC-NPC-iNeuron transcripts per million (TPM)

| Species | Tissue | NPEPPS_TPM | Total TBC1D3_TPM | Terminal_TBC1D3_TPM | CCL4L2_TPM | Fusion_TPM |
|---|---|---|---|---|---|---|
| Chimp | iPSC | 22.73 | 0 | 0 | 0 | 0 |
| Chimp | NPC | 8.61 | 0.1 | 0 | 5.7 | 5.3 |
| Chimp | neuron | 26.05 | 0.4 | 0 | 23.8 | 21.2 |
| Human | iPSC | 10.47 | 0.9 | 0.3 | 0 | 0.1 |
| Human | NPC | 30.03 | 6 | 2.7 | 0 | 2 |
| Human | neuron | 34.81 | 9.8 | 9.4 | 0 | 4.4 |
| Orang | iPSC | 16.71 | 19.4 | 0.8 | 0 | 0 |
| Orang | NPC | 16.54 | 10.1 | 0.3 | 0 | 0 |
| Orang | neuron | 28.36 | 52.7 | 0.5 | 0 | 0 |

**Supplementary Table 6.**
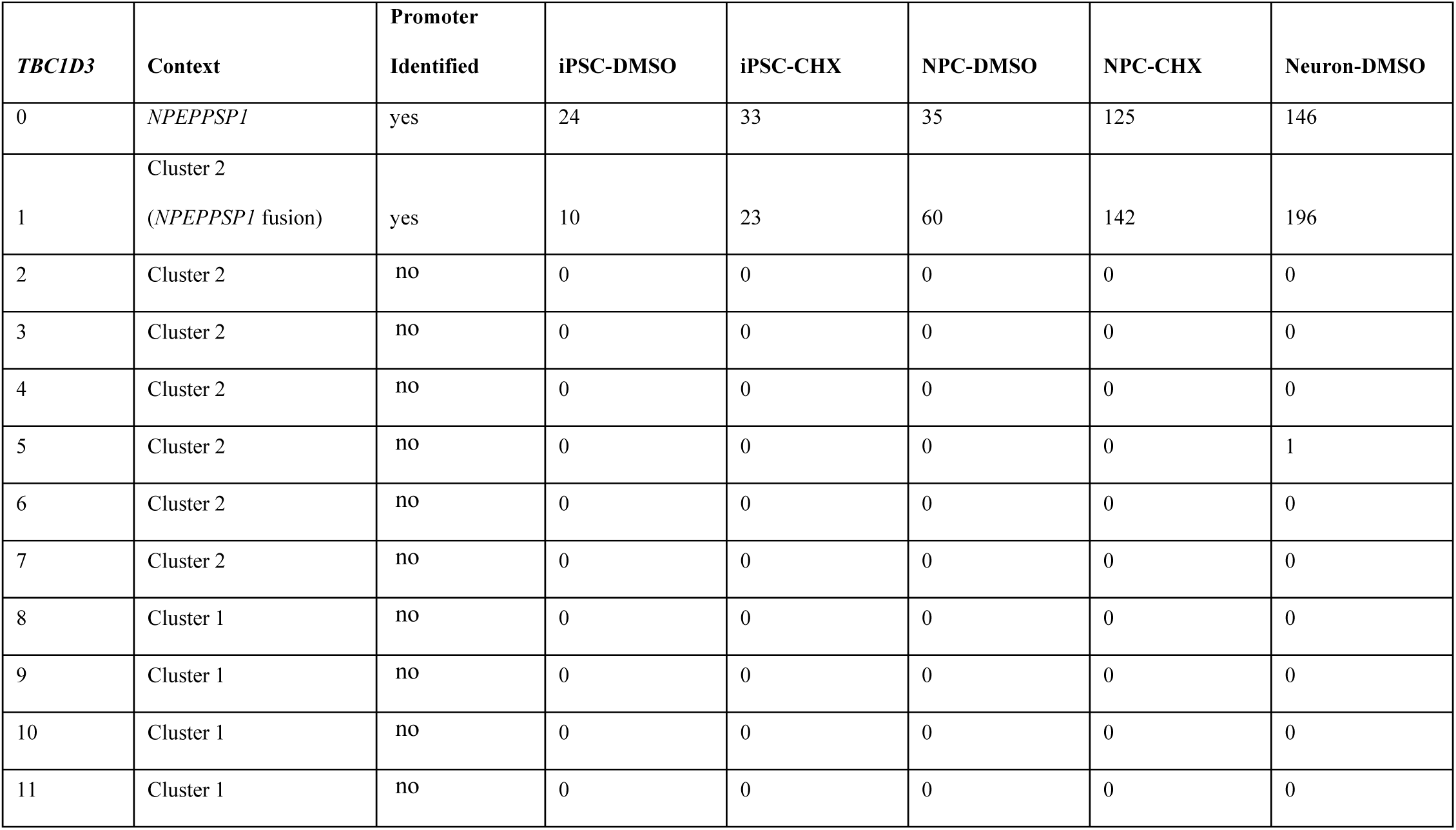
Human iPSC-NPC-iNeuron mappings by *TBC1D3* paralog.

| ***TBC1D3*** | **Context** | **Promoter Identified** | **iPSC-DMSO** | **iPSC-CHX** | **NPC-DMSO** | **NPC-CHX** | **Neuron-DMSO** |
|---|---|---|---|---|---|---|---|
| 0 | *NPEPPSP1* | yes | 24 | 33 | 35 | 125 | 146 |
| 1 | Cluster 2<br>(*NPEPPSP1* fusion) | yes | 10 | 23 | 60 | 142 | 196 |
| 2 | Cluster 2 | no | 0 | 0 | 0 | 0 | 0 |
| 3 | Cluster 2 | no | 0 | 0 | 0 | 0 | 0 |
| 4 | Cluster 2 | no | 0 | 0 | 0 | 0 | 0 |
| 5 | Cluster 2 | no | 0 | 0 | 0 | 0 | 1 |
| 6 | Cluster 2 | no | 0 | 0 | 0 | 0 | 0 |
| 7 | Cluster 2 | no | 0 | 0 | 0 | 0 | 0 |
| 8 | Cluster 1 | no | 0 | 0 | 0 | 0 | 0 |
| 9 | Cluster 1 | no | 0 | 0 | 0 | 0 | 0 |
| 10 | Cluster 1 | no | 0 | 0 | 0 | 0 | 0 |
| 11 | Cluster 1 | no | 0 | 0 | 0 | 0 | 0 |

**Supplementary Table 7.**
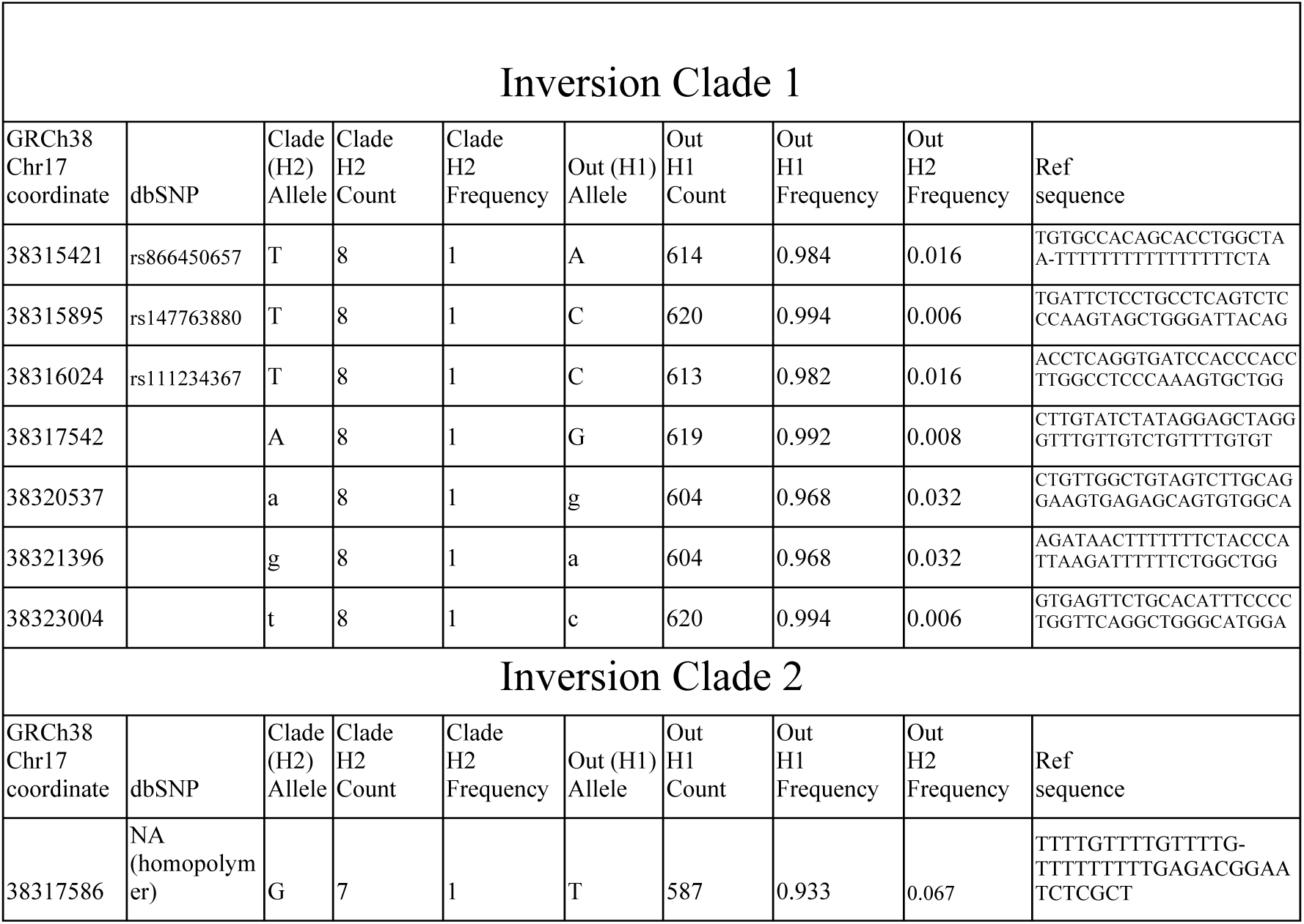
Inversion haplotypes by population.

**Supplementary Table 8.** Inversion Clade 1 tagging dbSNPs in UK Biobank.

| **UK Biobank dbSNP Genotype** |  |  |  |  |
|---|---|---|---|---|
| **rs147763880** | **Population** | **H1/H1** | **H1/H2** | **H2/H2** |
|  | Total | 402443 | 1156 | 31 |
|  | African | 5907 | 882 | 21 |
|  | East Asian | 1449 | 0 | 0 |
|  | Sout Asian | 6865 | 0 | 0 |
|  | European | 25583 | 8 | 0 |
| **rs111234367** | **Population** | **H1/H1** | **H1/H2** | **H2/H2** |
|  | Total | 401911 | 1623 | 89 |
|  | African | 5874 | 915 | 31 |
|  | East Asian | 1449 | 0 | 0 |
|  | South Asian | 6865 | 0 | 0 |
|  | European | 25583 | 8 | 0 |
| **rs112028599** | **Population** | **H1/H1** | **H1/H2** | **H2/H2** |
|  | Total | 402402 | 1190 | 89 |
|  | African | 5515 | 1226 | 69 |
|  | East Asian | 1449 | 0 | 0 |
|  | Sout Asian | 6865 | 0 | 0 |
|  | European | 25583 | 12 | 0 |

## METHODS

### Methylation Analysis

ONT BAM data used for the CHM13 methylation analysis were generated and processed described in (Mastrorosa et al., 2024). CHM13 gDNA reads used to generate the T2T-CHM13 reference were aligned back onto the reference using minimap2 (version 2.28; Li et al., 2018). Following alignment, methylation base calls of CpG sites were processed and aggregated using the modbam2bed software (v0.10.0; github.com/epi2me-labs/modbam2bed). Following methylation aggregation, CpG sites with coverage of five reads or less were discarded. For visualization (see Fig. 1), percent of reads methylated over each CpG site are illustrated as black points, with a rolling average of 15 CpG sites visualized with a red line. These results were visualized in R using ggplot2 geom_point and geom_line functions.

Similarly, methylation calling for ONT fast5 files from iPSCs, NPCs, and iNeurons, generated from human, chimpanzee, and orangutan, respectively, was generated during base calling by Dorado (version 0.8.2). These reads were then aligned to the respective species DSAs, characterized in (Jeong et al., in preparation) with minimap2 (version 2.28). Methylation calls were then aggregated over CpG sites with Modkit (version 0.4.1). The experimental design illustrated in Fig. 4A was generated with BioRender.com

### Fiber-seq Analysis

Fiber-seq libraries of HG02630 were prepared as described in (Real et al., 2025). Reads from the dataset were realigned to polished HG02630 assemblies generated with Deep polisher by the HPRC (Mastoras et al., 2025). Reads were mapped back to HG02630 with the following minimap2 (version 2.24), and adenosine and cytosine methylation calls from the original base calling were maintained with the new alignment BAMs. FIRE peaks were identified using FIREtools (version 0.0.7; Vollger, et al., 2025). Fiber-seq peaks were visualized with Gviz (version 1.46.1). The neurosphere experimental design illustrated in Fig. 2A was created with BioRender.com.

### Assembly Quality Control

To validate human and nonhuman primate assemblies, we investigated potential misassemblies over the *TBC1D3* Cluster 1–Cluster 2 locus with either NucFlag or HMM-Flagger pipelines (Gao et al., 2025; Liao et al., 2023). First, we required phased, contiguous assembly over the entire Cluster 1-2 locus for both sample haplotypes. Following this filtering, we intersected the Cluster 1-2 locus with QC BED files generated by both the HPRC and HGSVC. These included NucFlag (version 0.3.3) for 449 samples and HMM-Flagger (version 1.1.0) for 126 samples. We omitted assemblies that showed any indication of collapse, misjoin, duplication, or misassembly, allowing only “HAP” and “HET” annotations by the software pipelines.

### Full-Length Transcript Analysis

CDNA libraries were aligned as previously reported in (Guitart et al., 2024). Reads were aligned to a reference assembly using minimap (version 2.24) with the settings <minimap2 -ax splice --sam-hit-only --secondary=yes -p 0.5 --eqx -K 2G>. Following alignment, primary and secondary alignments were scrutinized. For paralog-specific reads, primary alignments with alignments scores greater than second-best alignments were used. Reads were visualized with R using the Gviz visualization library. Reads mapping to only a single exon were excluded from the gene transcript counts.

Iso-Seq libraries from multiple tissue sources were gathered as described in (Dishuck et al., 2025). Iso-Seq libraries were normalized to transcripts per million. Read counts were then organized by hierarchical clustering with R hclust and visualized as a heat plot with ggplot2.

### Polyadenylation Analysis

Polyadenylation signals were identified by looking for a canonical polyadenylation motif (AATAAA) within 20 bp of the end of transcript read alignments (Higgs et al., 1983). Splice junctions were identified by marking AG 3’ splice acceptor and GT 5’ splice acceptor motifs at splice junctions along cDNA alignments (Stephens & Schneider 1992). Polyadenylation signals and splice junctions were visualized with rtracklayer (version 1.62.0; Lawrence et al., 2009).

### Phylogenetic Analysis

Multiple sequence alignments (MSAs) for phylogenies were generated as follows. Homologous sequence was identified by mapping a reference target sequence to assemblies using minimap2 (-x asm20 --secondary --eqx -p -.05 -N 1000). Following sequence extraction, homologous sequences were aligned as an MSA with MAFFT (version 7.5.25; Katoh et al., 2013) with the following parameters: --reorder --preservecase --adjustdirection --maxiterate 1000 --thread 16. Following MSA generation, maximum likelihood phylogenies were generated with iqtree2 (version 2.1.2; Minh et al., 2020) with the following parameters: -nt AUTO -m MFP -s {MSA_fasta} -o {outgroup} --prefix {output_name} -alrt 1000 -bb 1000.

For *NPEPPSP1* duplicon phylogenetic analysis, the terminal 15 kbp of the *NPEPPSP1* duplication most proximal to *TBC1D3* was extracted using SEDEF annotations (Numanagic et al., 2018). The sequence was then mapped to the human, chimpanzee, gorilla, orangutan, and macaque telomere-to-telomere (T2T) assemblies using minimap2 with the default setting “-x asm20” (Yoo et al., 2025). These mappings were extracted with BEDTools and aligned into an MSA with MAFFT (version 7.525; Katoh et al., 2013). Next, the MSA was trimmed for gaps and misalignments with both Trimal (version v1.4.rev22) and manual pruning. Then, a phylogeny and timing estimates were generated using the iqtree2 software (version 2.12; Minh et al., 2020), with macaque as the outgroup. Timing estimates were calculated with iqtree2, using the following divergence times: (human–chimpanzee: 6.5 MYA; human–gorilla: 8.0 MYA; human–macaque: 24 MYA) as estimated in the fossil record (Dunsworth 2010; Stevens et al., 2013). The following command for timing estimation was used: iqtree2 -s {MSA} --date {divergence_date list} --date-ci 100 -te {phylogeny} --keep-ident -redo -o {outgroup_name} --date-tip 0 -alrt 1000 -bb 1000). For *CCL4L2*, the same process described as above was followed for the alignment and characterization of the *CCL4L2* gene model with 5 kbp of upstream sequence included.

### Short-Read-Depth Analysis

Short-read libraries from the 1000 Genomes Project (Sudmant et al., 2010). Illumina libraries were k-merized into 32 bp libraries with Meryl (version 1.4.1). Next, k-mer libraries were aligned to the T2T-CHM13 reference genome (Nurk et al., 2022) using FastCN (Pendleton et al., 2018), allowing for up to two mismatches between 32-mer and assembly alignments. Copy numbers over *TBC1D3*, *NPEPPSP1*, and the *NPEPPSP1* inferred promoter (Supplementary Fig. 12) were estimated by taking the average copy number over the given region, estimated by normalized read-depth coverage to unique diploid sequence across the genome.

### Conservation in Human Haplotypes

Gene sequences were aligned using an MSA with MAFFT (v7.525). Single-nucleotide variants (SNVs) and indels were identified by comparing each gene sequence to the reference gene sequence from T2T-CHM13v2.0. Variant functional consequences were annotated using VEP (v.111) with gene annotation (JHU RefSeqv110 + Liftoff v5.2; Shumate et al., 2021). Pathogenicity predictions for missense variants were obtained using AlphaMissense (Cheng et al., 2023), which were lifted over to T2T-CHM13 coordinates using UCSC LiftOver and complemented by PolyPhen and SIFT (version 20240502). Lollipop plots were generated using the R package trackViewer (v1.40.0).

### Haplotype Sequence Annotation

SDs for *NPEPPS-TBC1D3* haplotypes were annotated in the *de novo* assemblies with SEDEF (version 1.1; Numanagic et al., 2018). Duplicon tracks were generated for each assembly with DupMasker (Jiang et al., 2007) and run with the Rhodonite workflow (version 0.12; https://github.com/mrvollger/Rhodonite).

### RNA-seq Analysis

Short-read libraries from (Coorens et al., 2025) were used for the analysis. Reads were aligned to T2T-CHM13 using BWA (version 0.7.17), and transcript abundance was normalized using pydeseq2 (version 0.5.2). Read abundance for *NPEPPSP1-TBC1D3* fusion expression was specifically quantified by counting reads spanning the merged exon boundaries across the last and first exons of *NPEPPSP1* and *TBC1D3*, respectively.

### Reading Frame Maintenance

Gene sequences were aligned using an MSA with MAFFT (v7.525). SNVs and indels were identified by comparing each gene sequence to the reference gene sequence from T2T-CHM13v2.0. Variant functional consequences were annotated using VEP (v111; McLaren et al., 2016) with gene annotation (JHU RefSeqv110 + Liftoff v5.2). Pathogenicity predictions for missense variants were obtained using AlphaMissense (Cheng et al., 2023), which were lifted over to T2T-CHM13 coordinates using UCSC LiftOver and complemented by PolyPhen and SIFT (20240502). Lollipop plots were generated using the R package trackViewer (v1.40.0).

### Mass Spectrometry Analysis

Amino acid sequences for *NPEPPSP1* and *NPEPPS* were pulled from RefSeq annotations and aligned into an MSA with MAFFT (v7.525). Amino acid substitutions between the two paralogs were determined visually from the alignment. Next, tryptic peptides for both protein sequences were computationally inferred with the Expasy online tool (https://web.expasy.org/peptide_cutter/). Diverse peptide sequences between the two proteins were then selected and isolated for later analysis. These peptides were searched for in the MassIVE database (https://massive.ucsd.edu/ProteoSAFe/static/massive.jsp?redirect=auth).

## DATA ACCESS

CHM13 methylation data was previously described in Nurk et al., 2022. Prefrontal cortex Iso-Seq data mapped to T2T-CHM13 was previously reported in Leung et al., 2021, with data available on SRA under BioProject identification number PRJNA664117. Fiber-seq data was previously reported in Real et al., 2025, and deposited on SRA under BioProject identification number PRJNA1236375. HG02630 assemblies used for Fiber-seq analysis were previously reported in Mastoras et al., 2025. Iso-Seq and Kinnex libraries explored in Supplementary Fig. 17 were previously described in Dishuck et al., 2025. Primate assemblies investigated were generated as part of the T2T and primate T2T projects, reported in Nurk et al., 2022, and Yoo et al., 2025. *De novo* assemblies and transcriptomic sequence generated for human, chimpanzee, and orangutan iPSCs, NPCs, and iNeurons, were developed and reported in (Jeong et al., in preparation). Short-read gDNA data used for WSSD investigation was previously reported in Sudmant et al., 2010.

## ACKNOWLEDGEMENTS

Research reported in this publication was supported, in part, by the National Human Genome Research Institute of the National Institutes of Health (NIH) under Award Numbers R01HG002385 and R01HG010169 (to E.E.E.) and R01AG087959 and R01MH134981 (to A.A.P.). The content is solely the responsibility of the authors and does not necessarily represent the official views of the NIH. Additional funding was provided by Weill Neurohub (to E.E.E. and A.A.P.), the Pershing Square Foundation (to A.A.P.), and Schmidt Futures Foundation (to A.A.P.). This project was funded in part by the Emory National Primate Research Center Grant No. ORIP/OD P51OD011132. This research has been conducted using the UK Biobank Resource under application number 51663. A.A.P. is a New York Stem Cell Foundation Robertson Investigator. E.E.E. is an investigator of the Howard Hughes Medical Institute.

## AUTHOR CONTRIBUTIONS

X.G. and E.E.E. conceived the project; X.G. performed the analyses relevant to all manuscript figures; J.W.B. generated and assisted in analysis of data related to human, chimpanzee, and orangutan neuron development model; L.R. conducted synonymous, missense, and nonsense analysis illustrated in Supplementary Fig. 9B,C; H.J. and D.Y. assisted in analysis of human, chimpanzee, and orangutan neuron development model; D.Y. also assisted in characterization of chromosomal inversions described in Fig. 3C and Supplementary Figs. 4–6; N.K. assisted with characterization of inversion dbSNPs in UK Biobank; D.P. assisted in characterization of human haplotypes structural diversity; M.R.V. assisted in analysis of methylation characterization of CHM13 gDNA. K.Hoekzema sequenced all ONT data. K.M.M., K.A.S., and M.A. prepared and sequenced HiFi data used in neurospheres and the comparative neuronal developmental model analyses. K.Hoglin, R.M., and B.P assisted in cell culture and experimentation of the comparative neuronal developmental model. A.A.P. assisted in the conception of the comparative neuronal developmental model. X.G. and E.E.E. wrote the manuscript with input from all authors.

## DECLARATION OF USE OF AI-ASSISTED TECHNOLOGIES

During the preparation of this manuscript, the authors used ChatGPT (OpenAI) to assist with language editing, rephrasing, and improving the clarity and readability of text originally written by the authors. All AI-assisted revisions were reviewed and edited by the authors, who take full responsibility for the scientific content and conclusions of the manuscript.

